# MoDorado: Enhanced detection of tRNA modifications in nanopore sequencing by off-label use of modification callers

**DOI:** 10.1101/2025.02.18.638820

**Authors:** Franziskus N. M. Rübsam, Wang Liu-Wei, Yu Sun, Bhargesh Indravadan Patel, Wiep van der Toorn, Michael Piechotta, Christoph Dieterich, Max von Kleist, Ann E. Ehrenhofer-Murray

**Affiliations:** Institut für Biologie, Lebenswissenschaftliche Fakultät, Humboldt-Universität zu Berlin, 10115 Berlin, Germany; Systems Medicine of Infectious Disease (P5), Robert Koch Institute, Berlin, Germany; Department of Mathematics and Computer Science, Freie Universität Berlin, Berlin, Germany; International Max-Planck Research School ‘Biology and Computation’, Max-Planck Institute for Molecular Genetics, Berlin, Germany; Klaus Tschira Institute for Integrative Computational Cardiology, University Hospital Heidelberg, Heidelberg, Germany; Department of Internal Medicine III (Cardiology, Angiology, and Pneumology), University Hospital, Heidelberg, Germany; German Centre for Cardiovascular Research (DZHK)-Partner Site Heidelberg/Mannheim, Heidelberg, Germany

**Keywords:** Pus1, Deg1, Pus4, SQK-RNA004, Ψ22, Ψ8, epitranscriptomics

## Abstract

Rapid and accurate identification of tRNA modifications is crucial for understanding their role in protein translation and disease. However, their detection on tRNAs is challenging due to their high modification density. Recently, modification calling models for nanopore direct RNA sequencing became available for pseudouridine (Ψ), m^6^A, inosine and m^5^C, as part of the Dorado basecaller. Applying the Ψ model to tRNAs, we have mapped both known and novel Ψ sites in *Schizosaccharomyces pombe* and assigned the responsible pseudouridine synthetases. Furthermore, we have developed MoDorado, an algorithm to detect modifications beyond those used in model training (“off-label use”) by measuring prediction differences of pre-trained machine learning models. By leveraging the Ψ/m^6^A/inosine/m^5^C models, MoDorado detected seven additional modifications (ncm^5^U, mcm^5^U, mcm^5^s^2^U, m^7^G, queuosine, m^1^A, and i^6^A), thus generating a tRNA modification map of *S. pombe*. This work demonstrates the potential of pre-trained models in determining the intricate landscape of tRNA modifications.

## INTRODUCTION

Transfer RNAs (tRNAs) play an elementary role in protein translation in all living cells. The correct function of tRNAs strongly depends on the chemical modification of individual nucleotides in the tRNA, which regulate tRNA folding, structure, stability, aminoacylation and decoding ^1^. Indeed, approximately 100 different modifications are currently known in tRNAs, and each individual tRNA carries 5 to 15 modified sites ^2, 3^. Modification levels at particular sites can vary, and in some instances, are dynamically regulated depending on external conditions ^4^. Additionally, certain tRNA modifications are interdependent, an aspect that remains understudied ^5, 6^. Importantly, defects in tRNA modifications cause human diseases (termed “RNA modopathies”) such as mitochondrial diseases, neurological disorders and cancer ^7^. Therefore, having simple and cost-effective methods to measure tRNA modifications at single-molecule resolution is crucial for elucidating disease mechanisms and may have diagnostic potential.

In recent years, high-throughput methods for transcriptome-wide detection of RNA modifications have advanced significantly, utilizing techniques such as chemical derivatization or antibody-based approaches combined with high-throughput sequencing. However, many of these methods present challenges related to selectivity and efficiency ^8^. Hence, there is a need for a single, robust method to detect multiple modifications at the same time. A powerful approach for this purpose is to employ direct RNA sequencing (dRNA-seq) using nanopore sequencing from Oxford Nanopore Technologies (ONT), which has demonstrated accurate prediction of several types of modifications on a single mRNA transcript ^9–11^. Although dRNA-seq was originally developed for long read sequencing, the method has been adapted to sequence short tRNAs ^9, 12–17^. These adaptions involve ligating adaptors to the 5’ and 3’ ends of deacylated tRNAs, which are then threaded through membrane-embedded biological pores, where the nucleobases passing through a pore generate characteristic changes in the ionic currents across the pore. The distinct chemical properties of each nucleobase create unique signatures in the ionic current, allowing the inference of the sequence of the canonical A/C/G/U bases (termed “basecalling”). Modified bases can then be identified 1) through characteristic signal deviations from the canonical bases, and 2) systematic basecalling errors arising from such signal deviations ^18^.

In December 2023, ONT released a novel RNA-specific pore chemistry (SQK-RNA004, designated “RNA004” below), which showed accuracy improvement over previous iterations with the latest basecaller Dorado ^19^. Subsequently, ONT released pre-trained modification calling models for pseudouridine (Ψ) and N^6^-methyl-adenosine (m^6^A) in May 2024, and for 5-methyl-cytosine (m^5^C) and inosine in September 2024. These models were trained using synthetic RNA molecules containing modifications at specific positions within randomized sequence contexts (a strategy termed “randomers”). Due to their vast diversity of sequence context, these randomer datasets potentially represent the highest possible quality training data achievable for modification detection models. However, the performance of these models beyond standard usage on synthetic RNA oligos, for instance on heavily modified tRNAs *ex cellulo*, remains to be determined ^20^.

As a naturally occurring class of RNA, tRNAs represent an ideal test case for the limits of RNA modification detection, since the sites of modification and the modification machinery in many cases are well known. Here, we present the first application of pre-trained modification-specific machine learning models to modification identification in a tRNA context (for overview, see Figure 1). In a first step, we evaluated the Ψ caller for its ability to directly map Ψ on tRNAs from the fission yeast *Schizosaccharomyces pombe* (“on-label” usage, Fig. 1, lower left). The predicted Ψ positions were subsequently assigned to the pseudouridine synthetases (Pus) using basecalling error differences between wild-type (wt) strains and strains with deletions in the seven respective genes, which revealed two previously unknown Ψ sites in *S. pombe* (Ψ8 and Ψ22) and indicated the existence of modification co-regulation circuits. Together, the combination of direct Ψ prediction and basecalling error analysis allowed us to generate a near-complete eukaryotic Ψ map. Moreover, we developed MoDorado (a modification detection algorithm based on pre-trained modification calling models in Dorado), which leverages prediction score distributions of existing machine learning models, a feature that so far has not been exploited in existing detection methods. MoDorado enabled us to detect seven additional modifications not used in model training, a strategy we coined “off-label use” (Fig. 1, lower right). Altogether, our study provides a powerful approach for the enhanced detection of tRNA modifications.

**Fig. 1.**
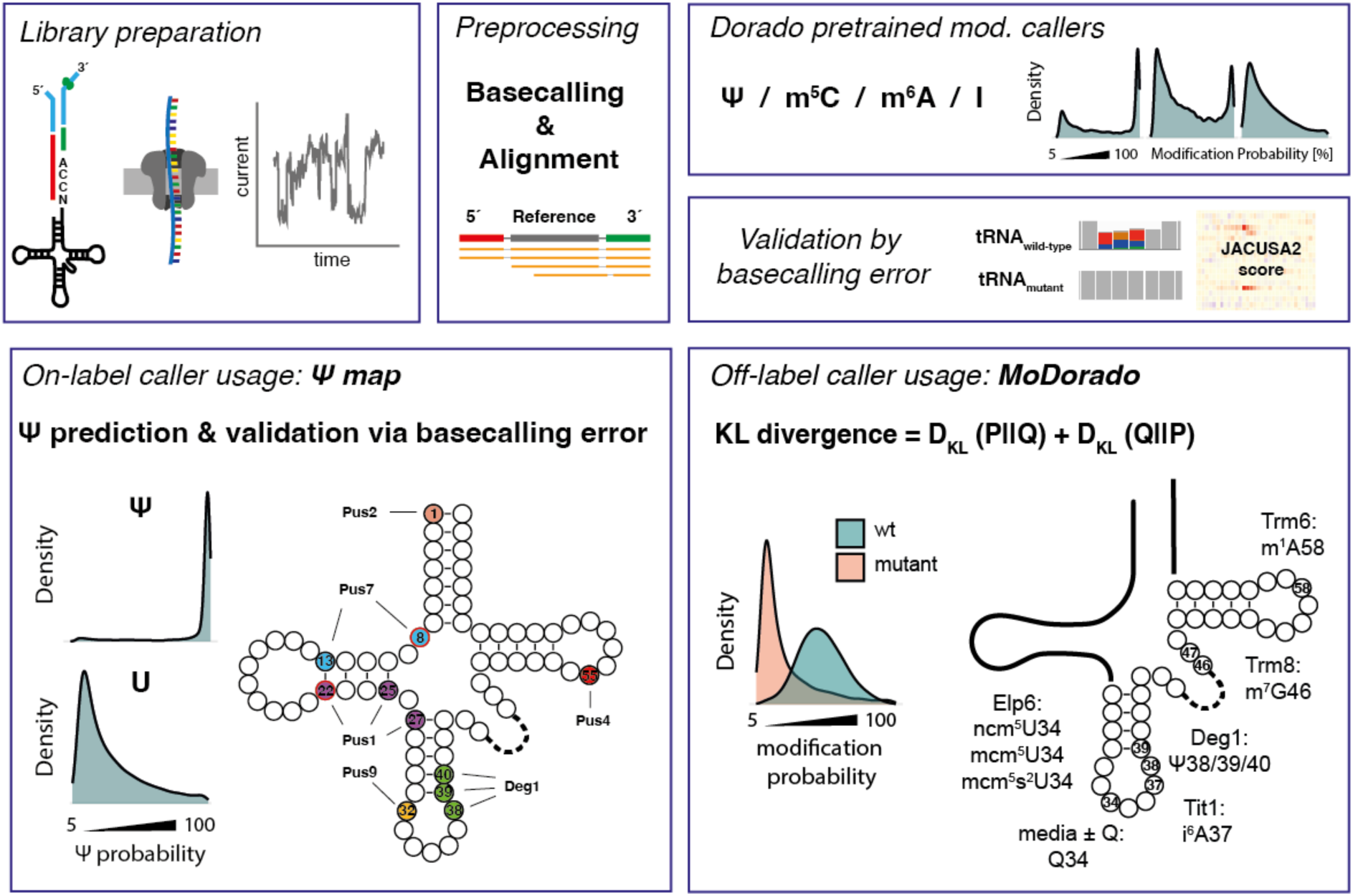
Strategy to determine tRNA modifications using nanopore dRNA-seq and MoDorado. For library preparation, splint adaptors were ligated to the 5’ and 3’ end of the deacylated tRNAs (top left). Raw data were basecalled with Dorado, and reads were aligned to the references using Parasail (top middle). The four pre-trained modification models for Ψ/m^6^A/inosine/m^5^C were used to predict modification probability for every nucleotide (plots show hypothetical examples of the distribution of probabilities). As validation of a modification, basecalling error differences between tRNAs from wild-type (wt) and strains lacking modification enzymes (mutant) were determined (top right). Using *pusΔ* strains, a near complete Ψ modification map was generated (“on-label usage”) and the responsible Ψ synthetases assigned to individual Ψ sites (bottom left). To detect modifications beyond those included for model training (“off-label usage”), we developed MoDorado, an algorithm that compares prediction score distributions of pre-trained machine learning models using the KL divergence. By comparing wild-type and gene deletion strains, MoDorado detected seven additional modifications: ncm^5^U, mcm^5^U, mcm^5^s^2^U, m^7^G, queuosine, m^1^A, and i^6^A (bottom right).

## RESULTS

### Direct Ψ calling in tRNAs reveals novel Ψ sites at positions 8 and 22

To investigate the utility of the Dorado Ψ caller on eukaryotic tRNAs, we isolated tRNAs from wild-type *S. pombe* and sequenced them using dRNA-seq (RNA004). Raw sequencing data was basecalled with Dorado using the Ψ model, followed by read alignment with Parasail ^21^ and filtering (see Methods). As expected, prediction probability distributions for sites annotated as Ψ-modified (Suppl. Table 1, Suppl. Fig. 1) generally tended to have a high peak near 100% probability, while non-Ψ sites tended to have a peak near 0% probability (Fig. 2A). For example, in Pro-TGG, Ψ55 ^22^ showed a unimodal distribution with a high peak near 100% Ψ probability, while the non-modified U11 had mostly low Ψ probability scores, which agrees with the lack of Ψ annotation (Fig. 2B). Interestingly, Ψ32 had a bimodal distribution, which may indicate partial modification ^23^, or may suggest that the prediction of Ψ is less accurate in certain sequence contexts. Two independent biological replicates showed high reproducibility (R^2^ = 0.984, Suppl. Fig. 2A).

**Fig. 2.**
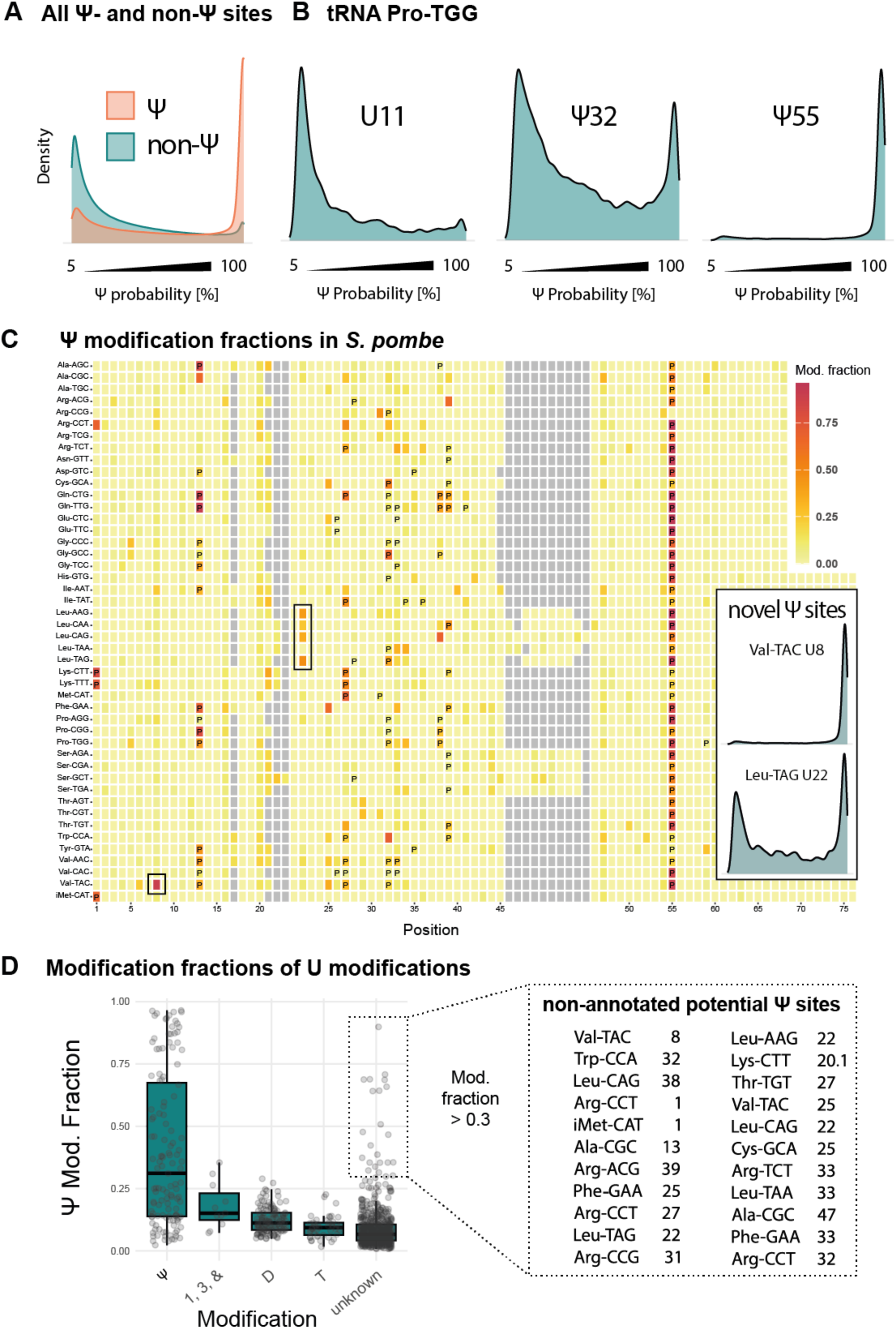
Determination of pseudouridine modification sites in tRNA from wild-type *S. pombe* reveals novel Ψ sites at positions 22 and 8. A) Distributions of prediction scores of all Ψ sites and sites that are not annotated as Ψ of all tRNAs (“non-Ψ”. 5% is the default minimum threshold in Dorado. B) Example distributions for annotated Ψ sites (32 and 55) and a non-Ψ site (U11) in tRNA Pro-TGG. C) Plot of modification fractions on individual *S. pombe* tRNAs. The x-axis denotes tRNA position; tRNAs are annotated on the y-axis. The colour indicates the modification fraction. Annotated Ψ sites are marked with P. Black boxes refer to inlets: Distribution of probability scores from Ψ basecalling for the so far unannotated positions U8 (top) and U22 (bottom). D) Bulky modifications at U34 and dihydrouridine (D) cause erroneously high Ψ fractions, and so far unannotated positions reveal likely Ψ modification. 1, mcm^5^U; 3, mcm^5^s^2^U; &, ncm^5^U. Non-annotated sites with high modification fraction (>0.3) are given on the right as potential non-annotated Ψ sites. The position 20.1 on Lys-CTT refers to an inserted position based on multiple sequence alignment.

The distinct distribution patterns of the Dorado Ψ caller at different sites highlight the need for a robust strategy to distinguish between Ψ-modified, non-modified, partially Ψ-modified, or otherwise modified sites. To summarize a single prediction distribution, we defined a “Ψ modification fraction” for each given position (see Methods) and plotted the fractions for each tRNA (Fig. 2C; each line represents a single tRNA, individual tRNAs are annotated on the y-axis). In general, the sites with high Ψ fractions were in good agreement with the annotated Ψ sites (Fig. 2C, D). Notably, there were also some previously non-annotated positions with a high Ψ fraction. One example is position 8 in Val-TAC (Fig. 2C, upper inlet), which has been described to be Ψ-modified in human embryonic stem cells ^24^, but not in yeast. Furthermore, position 22 in Leu-AAG, -CAA, -CAG and -TAG contained a high Ψ fraction (Fig. 2C, lower inlet; tRNA-Leu-TAA has A22 and therefore no Ψ22). To our knowledge, this is the first description of Ψ modification of position 22 in tRNA (see below for verification).

Other non-annotated tRNA sites with high Ψ fractions were observed at positions that are known to be Ψ-modified in other tRNAs (1, 13, 25, 31, 32, 38, 39, Fig. 2D). Notably, several U modifications other than Ψ resulted in elevated Ψ fractions with the Dorado Ψ basecaller, in particular dihydrouridine (D) and the complex U34 modifications 5-carbamoyl-methyluridine (ncm^5^U), 5-methoxycarbonylmethyluridine (mcm^5^U) and 5-methoxycarbonylmethyl-2-thiouridine (mcm^5^s^2^U), indicating that the Ψ model erroneously calls these modifications as Ψ. Altogether, the Dorado Ψ caller showed good correspondence with annotated Ψ sites and identified two new Ψ sites in *S. pombe* at positions 8 and 22.

### Assignment of predicted Ψ sites to known pseudouridine synthetases reveals Pus1-dependent Ψ22 and Pus7-dependent Ψ8

The above analysis showed that U modifications other than Ψ can produce signal changes that yield increased probability scores by the Ψ caller (Fig. 2D) ^25^. Therefore, to validate the predicted Ψ sites, we aimed to identify the pseudouridine synthetases responsible for their modification. In *S. pombe*, there are seven known tRNA Pus enzymes: the cytoplasmic tRNA Pus enzymes Pus1 ^26, 27^, Deg1 ^28^, Pus4 ^22^, Pus7 ^29^ and Pus9 ^23^, as well as the mitochondrial Pus2 and Pus3 (Suppl. Fig. 3) ^30^. To identify which enzymes modify which sites, we compared basecalling errors from dRNA-seq of tRNAs between wild-type and strains carrying deletions in the respective Pus genes using JACUSA2 ^18^. Ψ sites were assigned by defining an outlier threshold for the basecalling error scores (Suppl. Fig. 4 – 11, Suppl. Table 5, see Methods).

To directly call a site Ψ-modified or non-modified with the Dorado Ψ model, we searched for the optimal threshold for the Ψ fraction by plotting the percentage of annotated Ψ sites versus non-Ψ sites (Suppl. Fig. 2B). This showed a 63% recovery of Ψ sites and 8% of non-Ψ sites at a Ψ fraction of 0.2, which was used as a cut-off for Ψ detection *de novo* (i.e, with only the wild-type sample). Altogether, 82 sites called by the Dorado Ψ model could be assigned to a Pus enzyme based on the basecalling error score (Fig. 3A, B). 20 sites were assigned solely based on basecalling errors, but not by direct Ψ calling with Dorado. Importantly, the Ψ model and the error-based detection method draw from separate and complementary data sources: the Ψ model only outputs predictions for positions that were basecalled as U, whereas the error approach relies on reads in which U was miscalled (Fig. 3C). Lastly, 40 annotated sites, many located in the anticodon loop, were not detected by either Dorado or error profiling. This is likely due to the presence of many complex modifications in the close vicinity of Ψ sites (see below), or may be the result of mis-annotation.

**Fig. 3.**
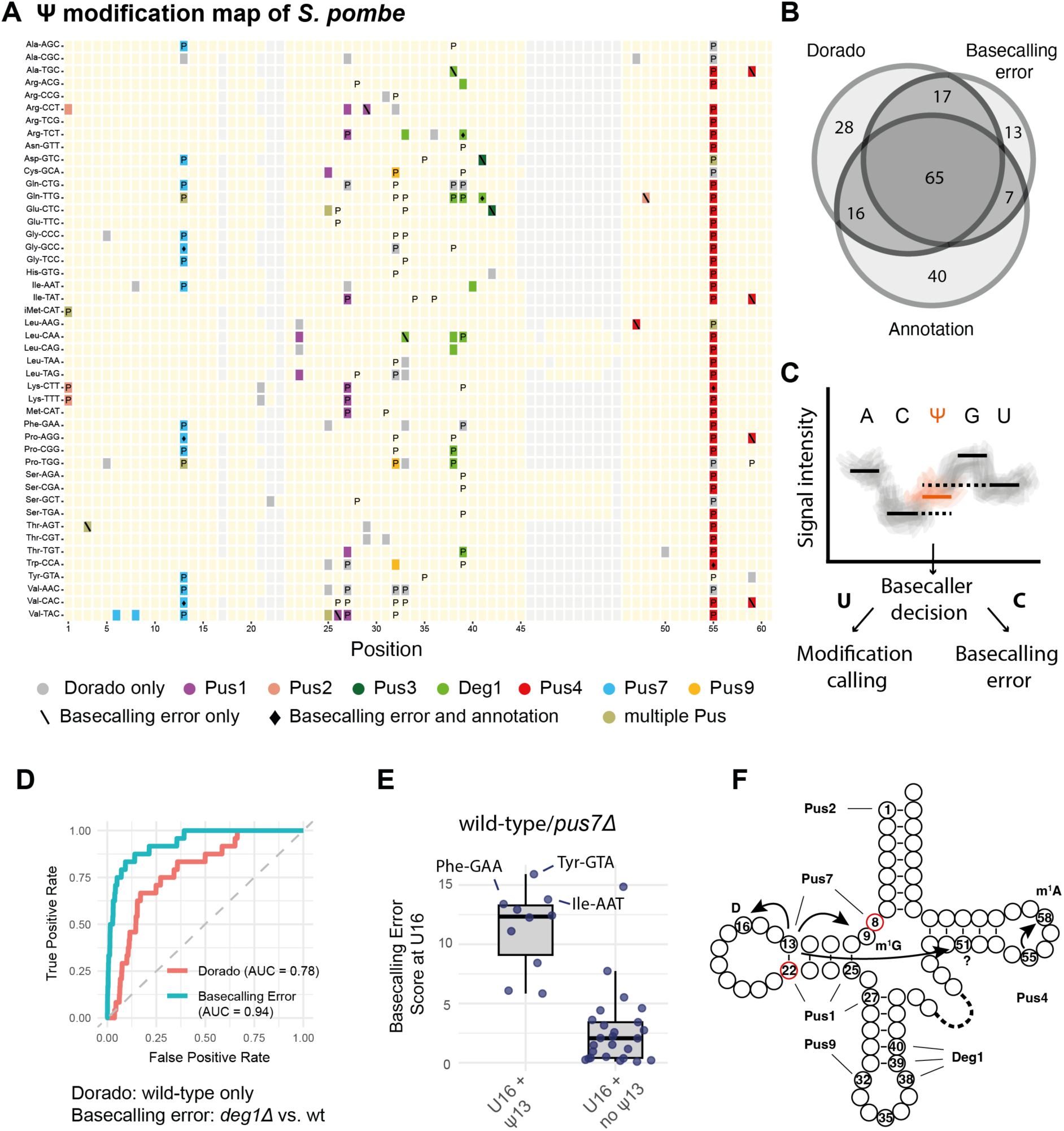
Determination of Ψ sites for pseudouridine synthetases and their tRNA modification circuits in *S. pombe*. A) Pus modification map in *S. pombe*. tRNAs are indicated on the y-axis; the x-axis denotes tRNA position. Annotated Ψ sites are marked with P. Ψ sites designated by direct Ψ basecalling and assigned to a pseudouridine synthetases based on basecallling errors are shown in colour. Sites that are only found by direct Ψ basecalling, but not assigned to an enzyme by basecalling errors, are shown in grey. Sites that are designated Ψ based on basecalling errors alone are marked with a backslash (**\**). Sites that are assigned by basecalling and annotated but not identified using direct Ψ basecalling are marked with a diamond (♦). B) Venn diagram showing the overlap between the Ψ sites from three categories: Dorado Ψ calling, basecalling error and annotation. C) Schematic of the theoretical signal space of a hypothetical 5-mer and the Dorado decision process. The basecaller Dorado first decides whether a base should be called as U, and only then, the Ψ caller will be applied to produce a prediction score. Thus, basecalling errors will not yield a corresponding Ψ prediction. D) ROC curves for direct Ψ calling with Dorado (from wt) and basecalling error (from wt/ *deg1Δ)* at Ψ positions 38, 39 and 40. E) Potential cross-talk between Pus7-dependent Ψ13 and D16. Elevated basecalling errors were observed at D16 in tRNAs with, but not without, Ψ13. F) Overview of Ψ modifications on cytoplasmic tRNA in *S. pombe* and assignment of pseudouridine synthetases from this study. The novel sites Ψ8 and Ψ22 are marked in red. Supernumerary circles depict positions 20.1 and 20.2. The potential modification crosstalk between Ψ13, D16, m^1^G9 and position 51 are shown with arrows. The known modification circuit between Ψ55 and m^1^A58 that was also detected here is indicated with an arrow. For modification crosstalk, see also the supplementary discussion and Suppl. Fig. 13.

An integrated view of the assignment of Ψ sites to Pus enzymes leads to the following conclusions (Fig. 3A, Suppl. Fig. 12): Pus7 modifies positions 6, 8 and 13 in *S. pombe*. For position 13 in the D arm, this is consistent with prior knowledge of Pus7 ^29^. Dependence on Ψ8 on Pus7 is consistent with the fact that U8 is in UN**<u>U</u>**AR, the consensus motif of Pus7 ^31^. Pus1 modifies positions 25 and 27 in the D arm of tRNAs ^26, 27^. The newly identified Ψ22 site in the D arm is modified by Pus1. Pus9 modifies position 32 in the anticodon loop ^23^. Deg1, which is the direct homolog of Pus3 from *S. cerevisiae* and humans, modifies positions 38, 39 and 40 in the anticodon stem-loop ^28^. Pus4 modifies U55 ^22^. Pus2 modifies position 1 in three tRNAs, and Pus3 modifies iMet-CAT at position 1. This is unexpected, as both enzymes are thought to only modify mitochondrial tRNAs.

To compare the performance of the Dorado Ψ model (*de novo* prediction from wt *S. pombe*) and the basecalling error approach (comparing wt strains with strains deleted for *pus* genes), we computed the Area under the Receiver Operating Characteristic (ROC-AUC) using *deg1Δ,* which lacks Ψ38, Ψ39 and Ψ40. The error approach had a higher AUC (0.94) than Dorado (0.78) (Fig. 3D), showing that *de novo* Ψ prediction remains challenging in tRNAs. In addition, the error approach can detect cross-modification crosstalks. As a point in case, we observed increased basecalling errors in wt compared to *pus4Δ* not only at the expected Ψ55 ^22^, but also at position 58 (Suppl. Fig. 8), which is N^1^-methyl-adenosine (m^1^A)-modified by Trm6/Trm61 ^32^, thus confirming the known modification Ψ55-m^1^A58 circuit ^33^. Similarly, we observed basecalling error patterns indicative of Pus7-dependent Ψ13 inhibiting Trm10-mediated m^1^G9 ^34^ and enhancing Dus1-dependent D16 modification ^35^ (Fig. 3E and Suppl. Fig. 13, see also supplementary discussion). Figure 3F provides a summary of all Pus sites and their associated crosstalk described in this study.

### Off-label use of pre-trained modification callers to detect other tRNA modifications

Our above analysis indicated that U modifications other than Ψ can produce distinctive signal signatures in nanopore sequencing that cannot be cleanly assigned by the Dorado Ψ caller to either U or Ψ (Fig. 2B, C). We reasoned that these distinctive modification signals may also create distinctive patterns in the model’s prediction scores, and thus, that the Ψ prediction scores might contain features that allow detection of these other modifications, even though the model was not specifically trained for their identification. This motivated us to develop MoDorado, a tool to detect diverse types of modifications beyond those supported by modification-specific Dorado models, a strategy that we coined “off-label use”.

To test this strategy, we performed dRNA-seq of tRNAs from an *elp6Δ* strain, which lacks the Elongator complex required for performing several U34 modifications (ncm^5^U, mcm^5^U and mcm^5^s^2^U) ^36, 37^. Raw signal intensities as well as Dorado Ψ probability scores showed clearly distinguishable patterns between wt and *elp6Δ* (Fig. 4A, B). To assess the (dis)similarity between probability score distributions of wt and mutant, we defined the symmetric Kullback-Leibler divergence (KL-divergence) as a distance metric for modification detection between samples, with high KL-divergence indicating the presence of modification (see Methods). Applying this to wt compared to *elp6Δ*, high KL-divergence values were observed at the expected U34 positions (Fig. 4C), demonstrating that the Dorado Ψ caller is sensitive to other uridine modifications. Comparing KL-divergence to basecalling error scores showed that these U34 modifications led to increased error scores with a strong spreading effect both up- and downstream, while the KL-divergence score was centred around U34 and its direct neighbouring bases (Fig. 4D).

**Fig. 4.**
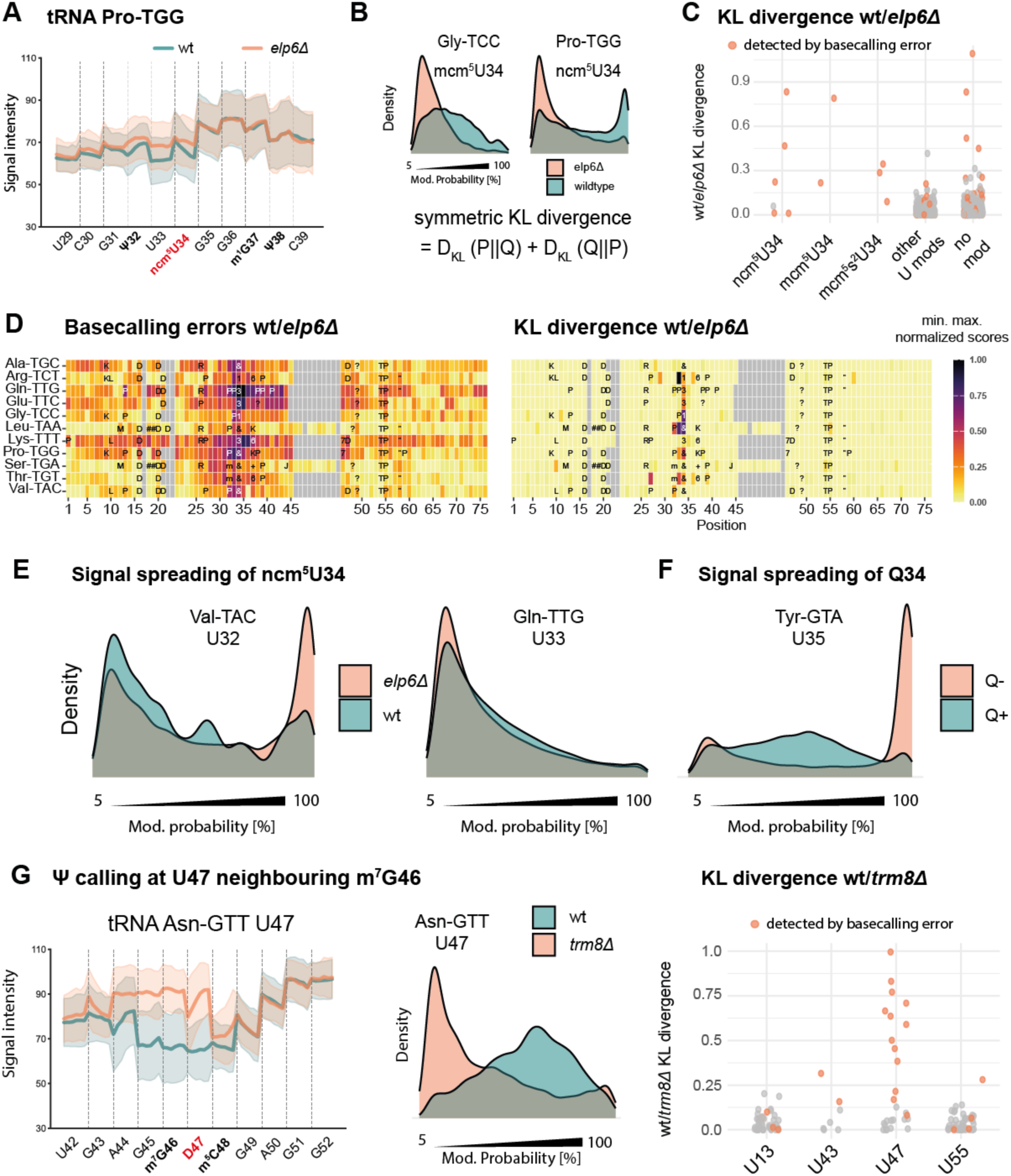
Off-label use of the Dorado Ψ caller to detect other U modifications (“MoDorado”), and detection of m^7^G by the Ψ caller at a neighbouring U. A) Raw signals for U29-C39 for tRNA-Pro-TGG from wt and *elp6Δ*, which carries ncm^5^U at position 34. Solid lines represent the mean signal intensity at a given position across reads. The shaded area represents one standard deviation of the signal intensity. B) Modification probability distributions, left: abnormal distribution for U34 in tRNA Gly-TCC, right: distribution for U34 in tRNA Pro-TGG. Bottom: definition of the symmetric KL divergence. C) KL divergence of all Elp6-dependent tRNA modifications, other U modifications, and non-annotated sites. Sites also detected by basecalling errors are shown in colour. D) Comparison of basecalling errors (left) and KL divergence scores (right) for tRNAs with Elp6-dependent U34 modification. Scores were min–max normalized for comparison. E) Prediction for Ψ32 in tRNA-Val-TAC in the presence (wild-type, wt) or absence (*elp6Δ*) of U34 modification (ncm^5^U). F) Prediction of Ψ35 in tRNA-Tyr-GTA in the presence (+Q) or absence (-Q) of Q34 modification. G) Detection of m^7^G47 based on Ψ probability at the neighbouring U47. Left: raw signals of tRNA Asn-GTT U42 – G52 from wt and *trm8Δ*. Middle: m^7^G46 leads to an abnormal distribution of Ψ prediction at U47. Right: KL divergence scores at U positions. U47 is next to m^7^G46. U positions with increased basecalling errors in wt/ *trm8Δ* are shown in colour.

Importantly, the elevated KL-divergence at the close neighbours of modified U34 indicates that modification signatures can influence the predictions at neighbouring nucleotides. To investigate this, we compared the Ψ probability scores at U32 of tRNA-Val-TAC, which is annotated as Ψ32 ^23^, but lies close to the Elongator-modified ncm^5^U34. Ψ32 was not detected in wt *S. pombe* by the Dorado Ψ caller, nor was it detectable by basecalling errors (Fig. 3A). In *elp6Δ*, however, the Ψ prediction scores showed a clear Ψ signature at position U32, demonstrating that ncm^5^U34 impairs direct Ψ calling at U32 (Fig. 4E, left panel). Similarly, a Ψ35 that lies close to queuosine 34 (Q34), was only identified in the absence of the queuosine modification (Fig. 4F), and Q34 caused erroneous Ψ calling at some U32 sites (Suppl. Fig. 14). These results are consistent with the notion that the reduced ability to detect Ψ in the anti-codon loop results from interference by other (bulky) modifications in the vicinity. Of note, some other annotated Ψ33 sites (e.g. Gln-TTG, Fig. 4E, right panel) showed no difference in prediction scores in *elp6Δ*, suggesting possible mis-annotation.

Currently, there is no Dorado model available for G modifications. Given the observation that current signals from one modification can spread into a neighbouring U position, we investigated whether off-label use of the Ψ model could be employed to detect G modifications that lie next to a U. We therefore sought to detect Trm8-dependent m^7^G46 ^38^ by computing the KL-divergence of Dorado Ψ probabilities at the neighbouring U47 (which is annotated as D47 ^35^) in wt and *trm8Δ*. Indeed, m^7^G caused signal distortion at and around position 46, which resulted in an elevated KL-divergence score at U47 (Fig. 4G), showing that U-neighbouring G modifications can be detected with “off-label use” of the Ψ model.

### MoDorado integrates multiple modification callers for a full tRNA modification map

Since the Dorado Ψ model exhibited broad sensitivity towards other U and neighbouring G modifications, we next evaluated the remaining three Dorado modification models (m^6^A, inosine and m^5^C) for their utility in detecting other tRNA modifications. Conveniently, since m^6^A itself is absent from tRNAs ^1^, all adenosine modifications detected by the m^6^A model by default constitute “off-label use” of the model. We investigated Tit1-dependent N^6^-isopentenylated A37 (i^6^A) ^39^ and Trm6/Trm61-dependent m^1^A58 ^32, 40^ by analysing dRNA-seq of tRNA from *tit1Δ* and *trm6Δ* strains. The m^6^A model resulted in elevated KL-divergence at i^6^A37 positions in wt/ *tit1Δ* (Fig. 5A) and at m^1^A58 in wt/ *trm6Δ* (Fig. 5B), thus establishing off-label use of the m^6^A model for the detection of both i^6^A and m^1^A.

**Fig. 5.**
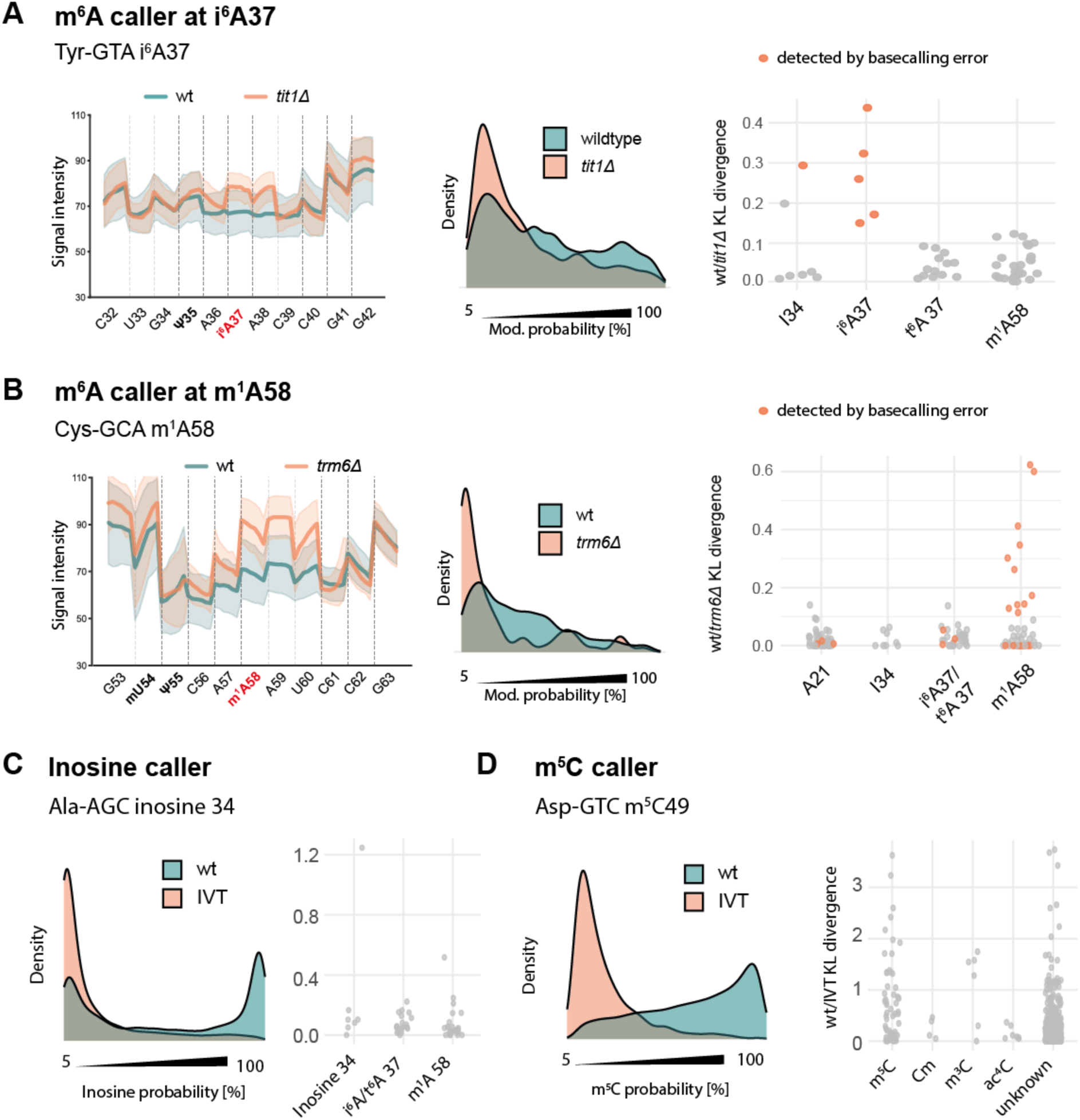
Evaluation of the m^6^A, inosine and m^5^C models on tRNAs. A) Off-label detection of i^6^A37 with the m^6^A model in wt/ *tit1Δ.* Left, raw signals of tRNA-Tyr-GTA C32 to G42 in wt and *tit1Δ*. Representation as in Fig. 4A. Middle, modification probability distributions generated by the m^6^A caller at A37 of tRNA-Tyr-GTA. Right, KL divergence at A37 and other A positions in wt/ *tit1Δ*. Positions with elevated basecalling errors are shown in colour. B) Off-label detection of m^1^A58 with the m^6^A model in wt/ *trm6Δ.* Left, raw signals; middle, modification probability distributions; right, KL divergence scores. Representation as in A. C) Evaluation of the inosine model. Left, inosine probability distributions from wt and *in vitro*-transcribed (IVT) tRNA-Ala-AGC inosine 34. Right, KL divergence scores at known inosines and other A positions. D) Evaluation of the m^5^C model. Left, m^5^C probability distribution at the known m^5^C49 position in tRNA-Asp-GTC. Right, KL divergence scores at known m^5^C sites and C position carrying other modifications.

Inosine and m^5^C are known to exist in several *S. pombe* tRNAs ^41–43^, thus allowing the evaluation of the Dorado inosine and m^5^C models. Since the adenosine deaminase Tad2/ Tad3 required for inosine formation is essential for viability in *S. pombe* ^42^, we created a library of *in vitro*-transcribed (IVT) tRNAs as unmodified tRNAs for determining KL-divergence scores. Furthermore, inosine modifications were verified by reverse transcription and Sanger sequencing, which confirmed their presence in all seven previously annotated tRNAs (Suppl. Fig. 15A). Interestingly, an increased KL-divergence score for inosine was only found at one of the seven known inosine sites (Fig. 5C). Other nearby modifications likely interfered with the detection, because inosine 34 in Ser-AGA was correctly detected in the absence of i^6^A at position 37 (*tit1Δ*, Suppl. Fig. 15B).

The levels of m^5^C in tRNAs from *S. pombe* are well known from earlier work ^44^. The KL-divergence scores were elevated at many of these sites (Fig. 5D), but m^5^C detection with the m^5^C model was not as reliable as Ψ with the Ψ model. Furthermore, m^3^C also showed elevated values, but since this modification is found in the anti-codon loop and we did not cross-verify the scores with the respective gene deletion or another orthogonal method, it is possible that the m^5^C model senses another adjacent modification, rather than m^3^C itself.

Having evaluated the four Dorado modification callers, we next combined them into a single MoDorado framework to assign KL-divergence scores to every A, U and C base in the tRNAs (no G model exists yet), and we used this approach to produce a dRNA-seq-based full tRNA modification map. To this end, tRNAs from wt strains were compared to IVT tRNAs using the m^6^A/Ψ/m^5^C models for A/U/C sites, respectively. Fig. 6A shows the normalized KL-divergence for each tRNA at single-nucleotide resolution. In general, higher KL-divergence was observed at modified sites compared to non-modified sites (e.g. Ψ or m^5^C), and as expected, the values were highest for on-label modifications. Multiple sites, especially A sites (e.g. position A23 in Ser-AGA and Thr-TGT or A15 in Arg-CCT and Lys-CTT), also showed high KL-divergence but, to the best of our knowledge, no modifications have previously been observed at those positions. An overview of KL-divergence scores of different modifications shows that the m^6^A/Ψ/m^5^C models sense a variety of modifications (Fig. 6B). In summary, MoDorado, which combines the on- and off-label usages of pre-trained modification callers, offers a promising approach towards determining the full tRNA modification map.

**Fig. 6.**
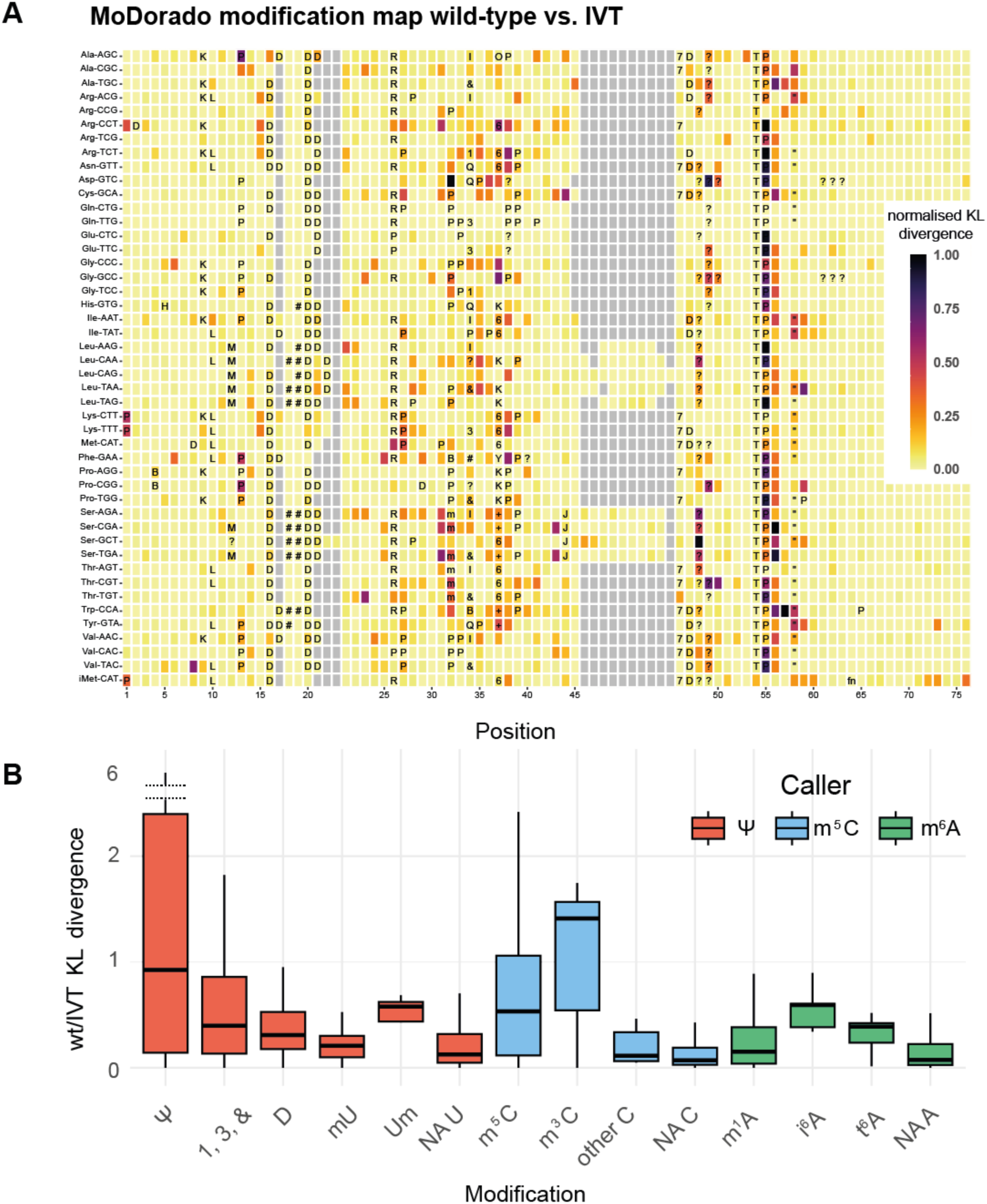
A: MoDorado integrates multiple models for a full tRNA modification map. A) Visualisation of KL-divergence scores between wild-type and IVT tRNAs in *S. pombe* using the Ψ, m^5^C and m^6^A models for U, C and A positions, respectively. Known and predicted modification sites (Suppl. Table 1) are marked using the unified RNAMods code for modified residues as in the Modomics database ^3^. B) KL divergence of various modifications in *S. pombe* tRNAs (wild-type/ IVT). Colours represent the nucleotide and the respective modification model used.

## DISCUSSION

The intricate network of modifications on tRNAs, often in close proximity, makes their detection and characterization a technically demanding task. Existing approaches for their detection using nanopore-based technologies generally use two types of features, namely, basecalling errors ^45^ or signal changes ^46^, or a combination of the two ^47^. In contrast, the MoDorado approach presented here uses as input the prediction scores of pre-trained modification-specific machine learning models, a feature that has not yet been exploited, but is readily available as part of basecalling with Dorado. This has allowed us to predict not only Ψ/inosine/m^5^C in tRNAs, but also other types of U/A/C modifications through the “off-label use” of these modification-specific callers, as well as G modifications by querying neighbouring bases. Altogether, MoDorado detected ncm^5^U, mcm^5^U, mcm^5^s^2^U, m^7^G, Q, m^1^A, i^6^A and (potentially) m^3^C modifications. In recent years, pre-trained models for m^6^A have shown promising results for the now retired RNA002 sequencing chemistry ^10, 48^. As more pre-trained models become available for the latest RNA004 chemistry and their performance continues to improve, we expect MoDorado’s capabilities to scale accordingly.

Among the modification callers available in Dorado, we found the Ψ caller to be the most consistent with previous Ψ annotations in tRNA, whereas the inosine and m^5^C callers showed lower consistency. Nevertheless, direct *de novo* modification calling remained challenging in the context of tRNAs, as is the case with the more traditional error-based approach with deletion mutants. This highlights the need for continued improvements of the modification-specific models and an integrative approach for detecting tRNA modifications.

Using a combination of methods, we have generated a tRNA pseudouridine modification map and have identified the respective pseudouridine synthetases, thus assigning all enzymes to their Ψ sites (Suppl. Fig. 19). This has led us to identify a novel target for Pus1, Ψ22 in four leucine tRNAs (tRNA-Leu-AAG, -CAA, -CAG and -TAG). This site lies opposite position 13 in the D arm of tRNAs, a position that is frequently Ψ-modified ^30^. However, the Ψ22-modified tRNAs have a G at position 13, suggesting that Ψ22 modification serves to enhance Wobble basepairing to G13 and thus to stabilize the tRNAs ^49^. We also identified a Pus7-dependent Ψ modification at position 8 in tRNA-Val-TAC, a site that so far has not been described in *S. pombe* ^24^.

In summary, the off-label use of pre-trained modification callers by MoDorado provides a novel and powerful framework for the interpretation of direct (t)RNA sequencing data using nanopore sequencing. With the release of alternative pre-trained models and their performance improvements in the near future, we anticipate that MoDorado will allow the comprehensive mapping of (t)RNA modifications, the identification of regulatory modification circuits, and the exploration of their impact on human health and disease.

## ONLINE METHODS

### *Schizosaccharomyces pombe* strain construction and growth

*S. pombe* strains used in this study are listed in Supplementary Table 2. Strains were grown in YES medium (5 g/l yeast extract, 30 g/l glucose, 250 mg/l adenine, 250 mg/l histidine, 250 mg/l leucine, 250 mg/l uracil, and 250 mg/l lysine) at 30°C. Synthetic queuine was supplemented to a final concentration of 0.1 µM as indicated. Strains from the Bioneer deletion collection were verified by amplification and sequencing of the bar code. Gene deletions of *deg1* (SPAC25B8.05), *pus4* (SPBC11C11.10), *pus7* (SPBC1A4.09),*tit1* (SPAC343.15) and *elp6* (SPBC3H7.10) were constructed using standard PCR-based integration of an antibiotic resistance cassette replacing the gene of interest. Deletions were confirmed by PCR. Primers used for the generation of gene deletions are listed in Supplementary Table 3.

### Extraction of small RNAs

Small RNAs were isolated as previously described ^13^. In short, *S. pombe* cells were cultured to an OD600 of 1, and 4 OD of cells were harvested by centrifugation. Cells were resuspended in 1 ml Trizol (Ambion) and 0.2 ml chloroform and glass beads were added. Cell lysis was performed by vortexing, followed by centrifugation at 16 000 rcf, 4°C for 15 min. After adding 215 μl ethanol to the upper phase, the sample was transferred to a spin cartridge (PureLink™ miRNA Isolation Kit (Invitrogen)) and centrifuged at 12 000 rcf for 1 min. 700 μl of ethanol was added to the flow-through, and the sample was transferred to a new spin cartridge followed by centrifugation at 12 000 rcf for 1 min. After washing the cartridge with wash buffer, small RNAs were eluted with 50 μl of DEPC-treated water. Small RNAs were deacylated for 30 min at 37°C in 100 mM Tris–HCl (pH 9) prior to library preparation.

### Detection of inosine by Sanger sequencing

For the detection of inosine, small RNAs where extracted from wt *S. pombe* cells as described above. tRNAs with annotated I34 (Ala-AGC, Arg-ACG, Ile-AAT, Leu-AAG, Ser-AGA, Thr-AGT and Val-AAC) were reverse transcribed with SuperScript IV (Thermo Fisher) using tRNA-specific RT primers (Suppl. Table 3), following the manufacturer’s protocol. Thereafter, the reverse transcription reaction was used as a template for cDNA amplification. Purified PCR fragments were subjected to Sanger sequencing, and inosine was detected by its characteristic A to G mutation (or T to C on the reverse strand, Suppl. Fig. 15).

### Generation of a library of representative *in vitro*-transcribed tRNAs from *S. pombe*

To obtain a mixture of representative, *in vitro*-transcribed tRNAs from *S. pombe*, a library of plasmids carrying the sequences of 46 *S. pombe* tRNAs was constructed. For this purpose, overlapping oligonucleotides (see Suppl. Tables 3 and 4) including a T7 promoter and a CCA tail were annealed, overhangs were filled in using Klenow polymerase and cloned into pJet1.2 using *Xho*I and *Nco*I (except *Sal*I/*Nco*I for Gln-tRNAs). Plasmids were amplified in *E. coli*, and equimolar amounts of the plasmids were pooled to generate the plasmid library.

To generate *in-vitro* transcribed (IVT) tRNAs, the plasmid library was digested using *Xho*I and *Nsi*I and purified by agarose gel extraction, which resulted in the release of a pool of 115 nt T7-tRNA-CAA IVT templates. 1 µg of template was used for IVT, which was performed at 37°C for 8 h using the TranscriptAid T7 High Yield Transcription Kit (Thermo Fisher Scientific). Samples were subsequently treated with DNaseI, and IVT tRNAs purified using phenol/chloroform/isoamylalcohol extraction followed by gel filtration on Sephadex G50 (GE Healthcare). tRNAs were treated with RppH (NEB) to generate 5’ monophosphorylated tRNAs, and samples were purified using the RNA clean & concentrate kit (Zymo Research).

### Library preparation for Nanopore Sequencing

In general, the preparation of tRNA libraries was carried out using the SQK-RNA002 or SQK-RNA004 kit (Oxford Nanopore Technologies, Suppl. Table 5) following the method previously described ^12, 13^ with minor modifications. Briefly, 100 pmol of deacylated small RNAs were refolded by heating, followed by ligation of the splint adapters using RNA ligase 2 (1x RNA ligase 2 buffer, 10% PEG8000, 2.5 mM ATP, 6.25 mM DTT, 6.25 mM MgCl2, 0.5 u/µl T4 RNA ligase 2, 20 pmol of each of the four splint adapters, 0.5 µl RNAse OUT) in a DNA loBind tube for 90 minutes at room temperature. Ligation reaction samples were subsequently separated on a 7 M urea/TBE PAGE gel (8%), and ligation products were excised and purified using the ‘crush and soak’ method. The concentration of gel-purified ligation products was measured with the Qubit RNA HS Assay Kit (Invitrogen). 500 ng of ligated tRNAs were used for library preparation according to the standard direct RNA sequencing ONT protocol for the respective kit (RNA002 and RNA004), and VATHS RNA Clean beads (Vazyme) were used for purification.

### Nanopore Sequencing

Samples were sequenced on a MinION Mk1C device using MinION FLO-MIN106D R9.4.1 or FLO-MIN004RA flow cells (Oxford Nanopore Technologies). After priming the flow cell, 75 μl of prepared sequencing library was loaded onto the flow cell. Sequencing runs were controlled by the MinKNOW software (Oxford Nanopore Technologies, version 22.10.5). Sequencing was typically carried out for 12 h depending on the number of collected reads and active pores. For FLO-MIN106D R9.4.1, runs of the raw bulk data file were stored, and the raw nanopore current intensity signals were re-processed with adjusted sequencing settings, as previously described ^16^. An overview of each dataset, the sequencing chemistry (RNA002 or RNA004) and other specifics are shown in Suppl. Table 5.

### Data processing

#### Reference curation

As reference sequences for *S. pombe* tRNAs, all unique mature tRNAs from GtRNAdb ^50^ and all unique mitochondrial tRNAs from Ensembl Fungi ^51^ were combined (total: 86 sequences), and the 5′ and 3′ splint adaptor sequences were added. Since mitochondrial tRNAs showed low sequence coverage, they were excluded from further analysis.

#### Annotation of pseudouridine and other tRNA modifications in S. pombe

Ψ annotations and other tRNA modifications are derived from the scientific literature and are either a direct demonstration of modification at a given site in *S. pombe*, or Ψ is inferred from *S. cerevisiae*. The individual sites and sources of information are given in Suppl. Table 1.

#### Basecalling and modification calling with Dorado

Base calling was performed using the official basecaller Dorado version rna002_70bps_hac@v3 (RNA002) and version rna004_130bps_sup@v5.1.0 (RNA004). For RNA004 data, the option to perform modification calling with the four currently available RNA modification models (Ψ/m6A/I/m5C) was used. For each individual read, the modification callers assign a discretized prediction score with integer value from 0 to 255 (representing a predicted probability of modification from 0 to 1; 0 to 100%) at positions of the corresponding canonical base (e.g. the Ψ model will only output scores for basecalled U positions). Only scores passing a threshold (default – 5%, which is discretized to a prediction score of 255 * 0.05 = 12) are written to output.

#### Read alignment and filtering

For alignment to the reference, all reads were taken and aligned to the reference using the optimal local sequence aligner Parasail (version 2.6.0) ^21^, as described ^13^. The output of Parasail are all-against-all read to reference alignments, which were filtered in the following steps: 1) for each read, only the reference(s) with the highest Alignment Score (“AS”) were included; 2) reads with AS less than 50 were excluded (threshold chosen according to the strategy described earlier ^13^); and 3) only full length reads were included.

### De novo detection of Ψ using the Dorado Ψ model

As basecalled reads contain errors, the positions with modification prediction scores may not actually exist (i.e. insertions), or certain reference positions may not exist in the reads (i.e. deletions). Therefore, only probability scores of correctly aligned (i.e. matched) positions were extracted (mismatched positions do not have a prediction score for the reference canonical base). Based on the prediction score distributions of Ψ-modified and non-modified U positions (Fig. 2A), a prediction score threshold of 204 (which is equal to 255 * 80%) was set. This threshold was used to define the fraction of reads with Ψ modification (N _reads with scores> 204_ / N _all reads_) for each U position.

### Modification detection from basecalling errors with JACUSA2

As an orthogonal method, JACUSA2 (version 2.0.4) ^18^ was used for modification detection based on basecalling errors on aligned reads. JACUSA2 outputs a single score per position, and modifications at positions with scores above the outlier detection threshold from the non-parametric Tukey’s fences method were determined:

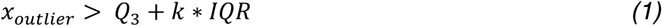

where Q_3_ is the third quartile, IQR is the interquartile range and k = 1.5. The resulting scores were further filtered (≥ 10) to facilitate data evaluation. The cut-off is based on the distribution of non-U base calling error outliers (Suppl. Fig. 11).

### Modification detection with MoDorado

The main objective of MoDorado is to leverage the output of existing pre-trained modification callers (e.g. Dorado) to detect modifications, by comparing the prediction score distributions between samples. The prediction scores of Dorado take discrete integer values from 12 to 255, which can be highly uneven in samples with lower coverage (a minimum coverage threshold of 100 per tRNA was used). Therefore, the score distributions were first smoothed by constructing frequency histograms with a fixed number of bins (n_bins = 10) of equal spacing between 11 and 256. To compare the similarity of the smoothed prediction score distributions between samples, the Kullback-Leibler (KL) Divergence for discrete probability distributions was used, which is given by equation 2.

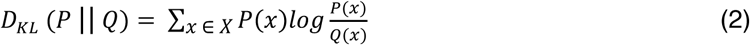

where P and Q represent the smoothed histogram distributions of the wild-type and the mutant samples, respectively. As the KL divergence are non-symmetric, the order of input distributions *P* and *Q* affect *D_KL_*. In MoDorado, the sum of *D_KL_* (*P* || *Q*) and *D_KL_* (*Q* || *P*) as a symmetric version of the KL divergence was used. Specifically,

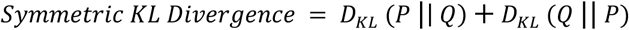

### Signal visualisation

During basecalling, the option `--emit-moves` was turned on to have Dorado output a signal segmentation (commonly known as the “move table”). The move table segmentation assigns each base of a read to a segment of signals stored in the pod5 files. Since basecalled reads contain errors, such as insertions and deletions, only the signals of aligned reference positions (matches and mismatches) were extracted for visualisation. As signals are variable in length (i.e., dwell time) and the signal lengths per base are always multiples of 6 (the fixed “stride” parameter in the RNA004 basecalling model), a subsample of 6 data points with equal spacing were taken for each base position. Lastly, to offset the intrinsic signal differences between two samples, we computed for each tRNA, 1) the differences in median signal intensity for each position between samples, and then 2) the median of these differences across all positions on the tRNA, which resulted in a single scaler number and was used as a vertical shift parameter to add to the signals of the mutant sample.

## Data availability

All raw sequencing data will be available on European Nucleotide Archive upon acceptance.

## Code availability

The analysis scripts for the paper and the source code for the MoDorado tool are available at https://github.com/KleistLab/MoDorado.

## Acknowledgments

We thank Eric Phizicky for providing *S. pombe* strains, Josta Hamann for technical support, and all lab members for insights and discussions. This work was supported by Deutsche Forschungsgemeinschaft (grants EH237/19-1 and 237/21-1 to A. E. E.-M.), Deutscher Akademischer Austauschdienst DAAD to B. I. P. and the Chinese Scholarship Council to Y. S. W.L.-W. was funded by the European Union’s Horizon 2020 research and innovation program under the MarieSklowska-Curie Actions Innovative Training Networks grant, agreement no. 955974 (VIROINF). M.v.K and W.L.-W. acknowledge funding provided by the German Ministry for Science and Education, grant number 01KI2016. M.v.K. acknowledges funding by the Deutsche Forschungsgemeinschaft (DFG, German Research Foundation) under Germanýs Excellence Strategy – The Berlin Mathematics Research Center MATH+ (EXC-2046/1, project ID: 390685689). C.D. and M.P. acknowledge funding by the German RMaP consortium (Deutsche Forschungsgemeinschaft [DFG, German Research Foundation, 439669440 TRR319 RMaP TP B01]).

## Author contributions

F. R., B. I. P. and Y. S. performed the wet lab experiments. F. R., W. L.-W. and W. vdT., C. D. and M. P. performed the bioinformatic analysis. A.E.E.-M. conceived the work. C. D., M. vK. and A.E.E.-M. supervised the work. A. E. E.-M., C.D. and M. vK. acquired funding to conduct the work. F. R., W. L.-W. and A. E. E.-M. prepared the figures. A. E. E.-M., F. R. and W. L.-W. wrote the paper, with contributions from all authors.

