## Supplementary Information for "MoDorado: Enhanced detection of tRNA modifications in nanopore sequencing by off-label use of modification callers"

### Supplementary Discussion

#### Site specificity of pseudouridine synthetases in *S. pombe*

We have employed dRNA-seq from wt and *pusΔ* strains to assign Pus enzymes to Ψ sites that were either determined by direct Ψ basecalling, annotated in the literature, or showed high basecalling errors in the wt/ *pusΔ* comparison and thus have been able to assign many Ψ sites to a particular Pus enzyme (see main text). Interestingly, some Ψ positions show high basecalling errors for more than one Pus enzyme. For instance, position 25 shows high error scores for Pus1 and Pus7 in several tRNAs and for Pus4 and Pus3 in single tRNAs (Fig. 3A). Since Pus7 mainly is the pseudouridine synthetase for Ψ13 [1], the concomitant change of error rate at Ψ25 could reflect a crosstalk between Pus7-dependent Ψ13 and Pus1-dependent Ψ25. Alternatively, Pus7 could directly Ψ-modifiy position 25 on some tRNAs.

Interestingly, position 1 also showed high basecallling error scores in several *pusΔ* strains, with most high scores in *pus2Δ*. Of note, Pus2 in a mitochondrial enzyme in *S. cerevisiae* [2], though its localization in *S. pombe* has not been determined. Perhaps *S. pombe* Pus2 has both cytoplasmic and mitochondrial functions. The wt/ *pus2Δ* dataset also showed elevated basecalling errors at position 13. Given the current knowledge about Pus2, the most likely explanation for this is a technical artefact.

The wt/ *pus3Δ* dataset showed very few positions with elevated basecalling errors. Perhaps Pus3 is a mitochondrial tRNA pseudouridine synthetase, or Pus3 modified RNAs other than tRNAs (for instance, mRNAs or snRNAs).

#### Pus7-dependent tRNA modification circuits between Ψ13, $m^1$G9 and D16

In this study, we employed the analysis of basecalling errors at non-U sites of tRNAs to identify potential cross-modification crosstalk. Interestingly, wt/ *pus7Δ* showed high basecalling errors at G9, U16 and G51 in several tRNAs (Fig. 3E, Suppl. Fig. 9, 13). Specifically, tRNAs that contain both G9 and Ψ13 showed high error scores at G9, whereas tRNAs with G9 but no Ψ13 showed low basecalling error scores at G9 (Suppl. Fig. 13A, left). Position G9 carries Trm10-dependent $m^1$G modification [3]. This indicates that the absence of Ψ13 in *pus7Δ* causes a change in the $m^1$G level at G9. Of note, the determination of basecalling error scores by JACUSA2 returns only positive values and therefore does not distinguish between high errors in wt or high errors in *pus7Δ* [4]. The observation that the absolute rate of mismatches (but not deletions or insertions) at position G9 is lower in wt compared to *pus7Δ* (Suppl. Fig 13A, right) suggests that the $m^1$G level is increased in *pus7Δ*. This indicates that there is a negative correlation between Pus7-dependent Ψ13 and Trm10-dependent $m^1$G modification. Of note, tRNA-Arg and tRNA-Glu also showed high basecalling error scores at position G9, but they carry C13, not Ψ13 (Suppl. Fig. 13A, red box).

An alternative explanation for higher basecalling error rates at a position other than the known modified site itself (in this case Ψ13) is that the signal from the modified base "bleeds" into surrounding regions and disturbs the basecaller at identifying the standard bases in the sequence close to the modification. We cannot exclude this possibility for Ψ13 and $m^1$G9. However, we note that basecalling errors are low at positions between G9 and U13, and we therefore favor the interpretation that the higher error rates are indicative of cross-modification crosstalk between Ψ13 and $m^1$G9.

We furthermore observed increased basecalling error scores in wt/ *pus7Δ* at position U16, which is modified to dihydrouridine (D16) by the enzyme Dus1 [5]. Furthermore, the mismatch error rates at position D16 were high in tRNAs with Ψ13, but not tRNAs that lack this position. Here, mismatch differences were higher in wt than *pus7Δ*, showing a positive correlation between Ψ13 and D16 and suggesting that the Ψ modification by Pus7 at position 13 enhances Dus1-dependent formation of D16 (Suppl. Figure 13B). Again, we cannot exclude the possibility that the higher error rates result from a spill-over of the signal of the Ψ base to surrounding sequences. Again, however, error rates were low in the intervening sequences, favoring the interpretation of a modification circuit.

Finally, two tRNAs (tRNA-Asp-GTC and -Gly-GCC) showed high basecalling error scores at G51 in in wt/ *pus7Δ* (Suppl. Fig. 13C). So far, no modification is known at position G51. Either this site carries a modification that remains to be detected and that shows cross-talk to Ψ13, or the increased error rate is a technical artefact of unknown source.

**Pus4-dependent tRNA modification circuit between Ψ55, T54 and $m^1A58$**

The Pus4 enzyme introduces Ψ at position 55, a site that is modified in nearly all tRNAs and found in all kingdoms of life, the human homologs being TRUB1 and TRUB2 (Suppl. Fig. 3) [6, 7]. In the wt/ *pus4Δ* dataset, we observed increased levels of basecalling errors at position 55 for 42 ot 44 tRNAs with annotated Ψ55 (Suppl. Fig. 8). The two remaining tRNAs (Ala-CGC, Cys-CGA) had low read numbers and therefore were not included in the analysis. Taken together, our data indicate that all tRNAs with U55 are Ψ-modified by Pus4 at this site, which is in agreement with earlier observations [8].

Two modifications in the T-arm (T54 and $m^1A58$) have previously been shown to be dependent on Ψ55 modification [9, 10], and this interdependency has been verified by dRNA-seq [11]. In agreement with these studies, we also observed high basecalling errors in the wt/ *pus4Δ* data in many tRNAs at positions 57, 58 and 59 (Suppl. Fig. 8). The increased error rate at A58 results from a decrease of $m^1A58$ in *pus4Δ* cells, whereas increased error rates at the surrounding positions 57 and 59 most likely result from spill-over of the $m^1A58$ signal to its neighboring positions. $m^1A58$ depends on the Trm6/ Trm61 enzyme [12]. Therefore, to investigate neighborhood effects of $m^1A58$, we performed dRNA-seq of tRNAs from *trm6Δ* and determined error rates compared to wt as described (see main text, Suppl. Fig. 16). Indeed, high basecalling error scores were observed at as well as around position 58, showing that error rates at positions 57 and 59 result from signal bleeding from $m^1A58$. Comparing the basecalling errors at position 57 and 58 in both datasets (wt/ *pus4Δ* and wt/ *trm6Δ*) suggests that the majority of tRNAs in *S. pombe* are $m^1A58$-modified. Moreover, our data identified more isoacceptors to be $m^1A58$-modified than found before in an earlier study [11]. This might be attributed to differences in cultivation conditions, because $m^1A58$ levels vary depending on the timepoint time point of harvesting the cells [11].

**Potential Deg1-dependent tRNA modification circuits**

Inspection of basecalling errors generated by Ψ shows that increased error rates are found at the Ψ position itself, but no or very little errors at neighboring positions, suggesting that signal distortions by Ψ only rarely spill over to surrounding sequences. It therefore was surprising that in wt/ *deg1Δ*, the basecalling error spreads extensively to the neighboring bases, sometimes from Ψ38 up to position 34 (examples are tRNA Arg-AGC,

Lys-CTT or Pro-TGG). Also position 37 frequently has high basecalling errors, a site that is known to carry several other modifications ($i^6A$, $t^6A$, $m^1G$). We therefore speculate that modification interdependencies might exist between Deg1-dependent Ψ38/ Ψ39 and modifications at position 34 or 37. However, this interpretation must be taken with caution and awaits independent verification using an orthogonal method.

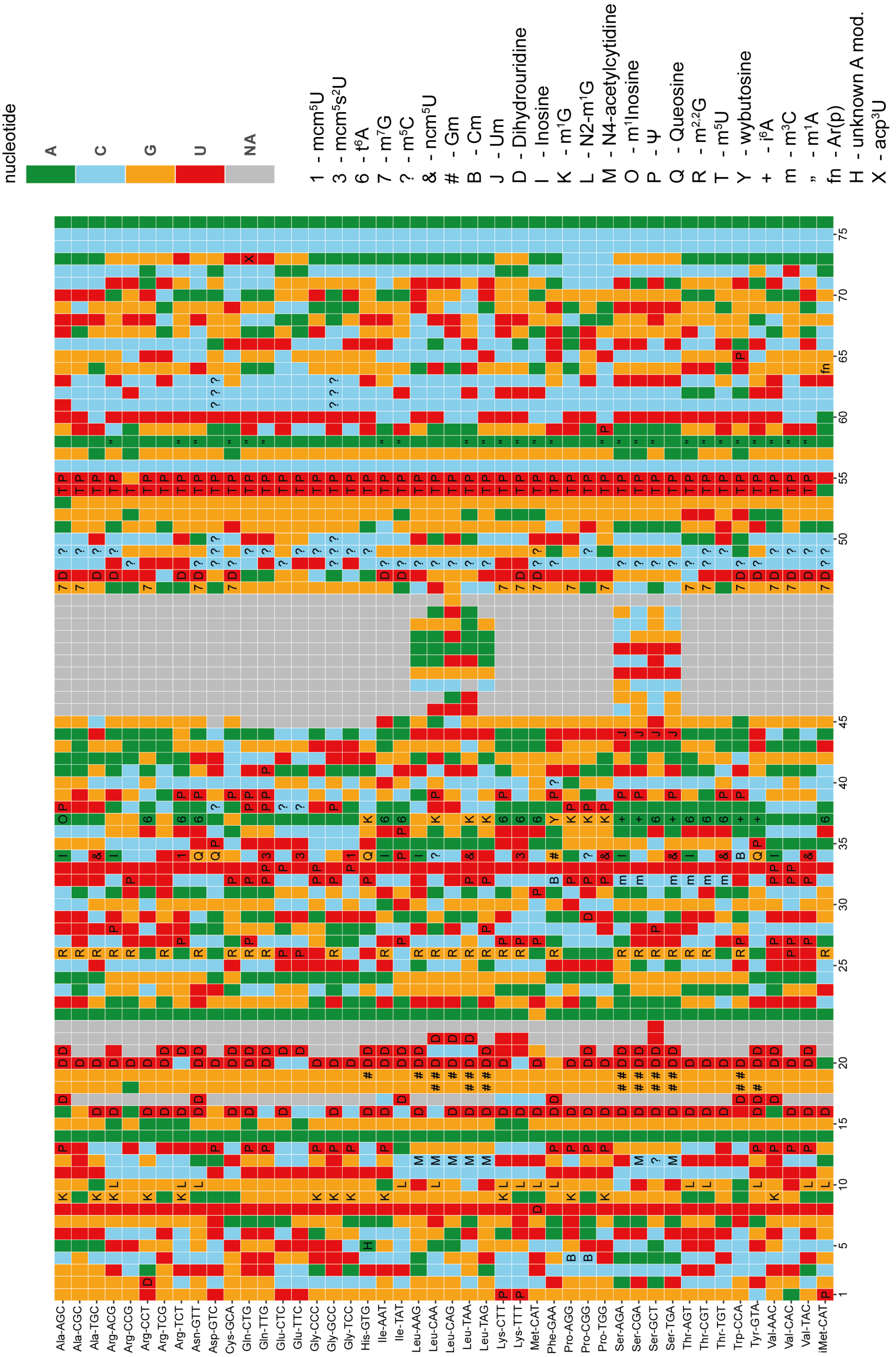


**Supplementary Figure 1**: tRNA modification map of *S. pombe* inferred from the literature or from *S. cerevisiae*. Sources are given in Supplementary Table 1. The modification annotation follows the MODOMICS one letter code [13]. Colour represents the reference nucleotide in the given tRNA at the given position: A = green, U = red, C = blue and G = orange.

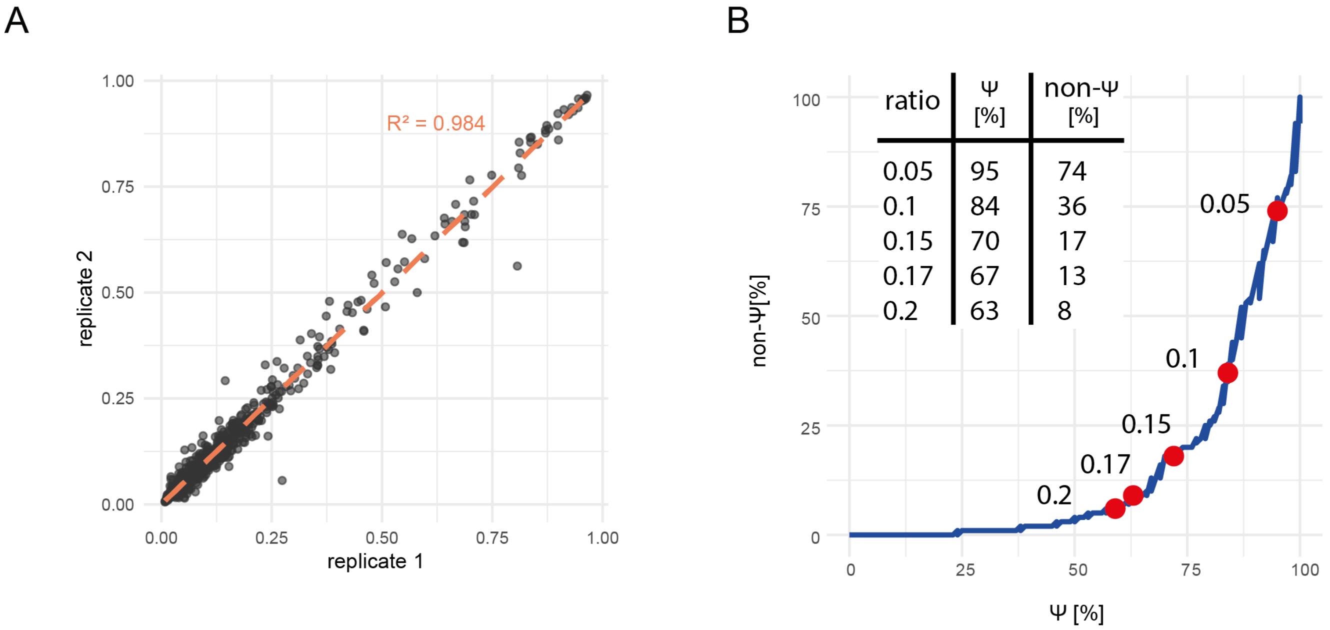


**Supplementary Figure 2**: A) Scatterplot of Dorado Ψ modification fractions between replicate 1 and replicate 2 of direct sequencing of tRNAs from wild-type *S. pombe*. B) Recovery rate of annotated and non-annotated Ψ sites at different modification fractions.

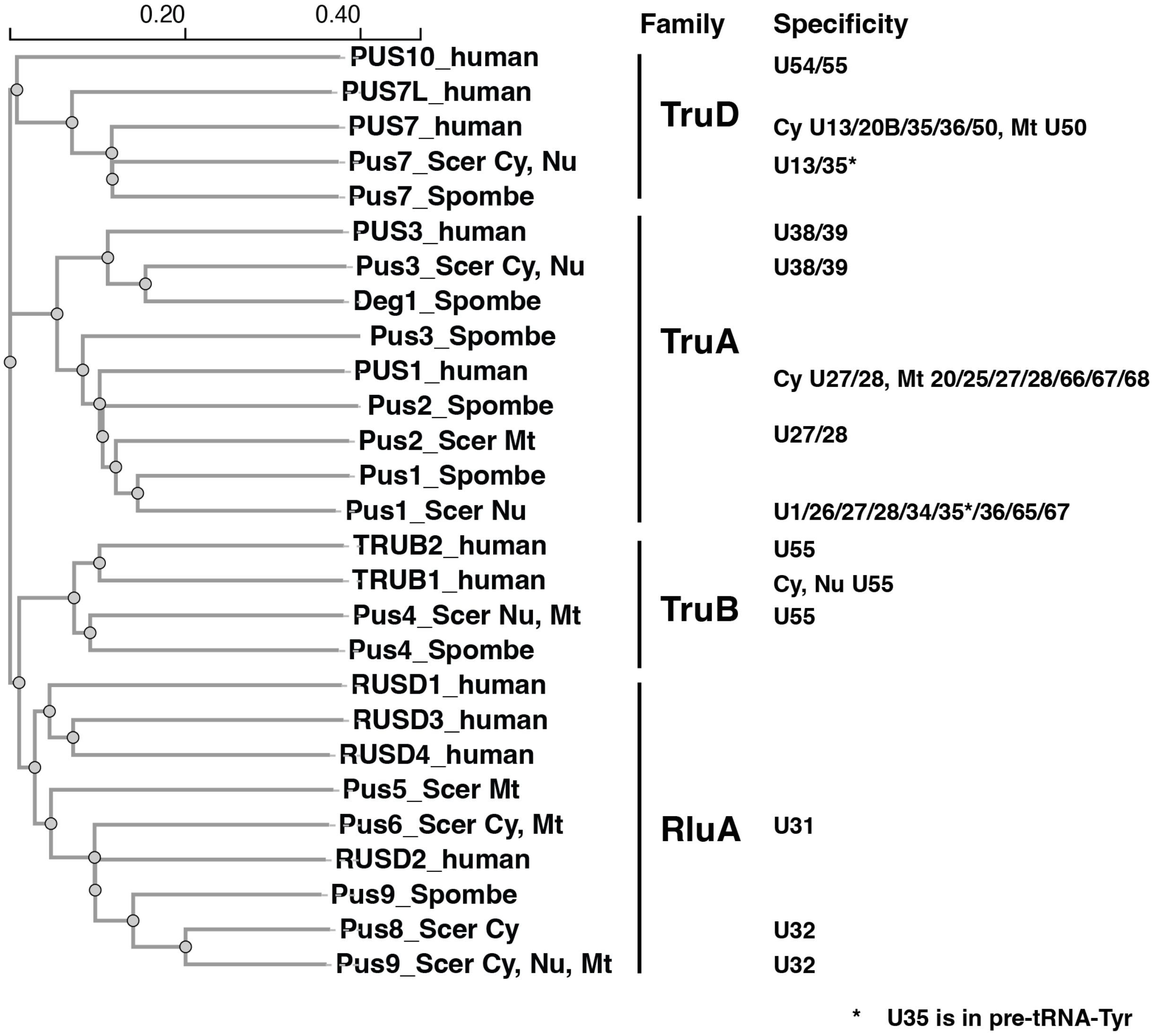


**Supplementary Figure 3**: Phylogenetic tree of pseudouridine synthases from *H. sapiens*, *S. cerevisiae* and *S. pombe*. The cellular localization and site specificity is noted where known (taken from [2]; Cy, cytoplasmic; Nu, nuclear; Mt, mitochondrial). The tree was generated by aligning protein sequences using Clustal Omega multiple sequence alignment using ClustalW and standard settings [14]. The following UniProt entries were used: O59721, O74343, O74451, O94295, O94396, O95900, P31115, P48567, P53167, P53294, Q06244, Q08647, Q09709, Q12069, Q12211, Q12362, Q3MIT2, Q6P087, Q8IZ73, Q8WWH5, Q96CM3, Q96PZ0, Q9BZE2, Q9H0K6, Q9UJJ7, Q9UTB3, Q9Y606

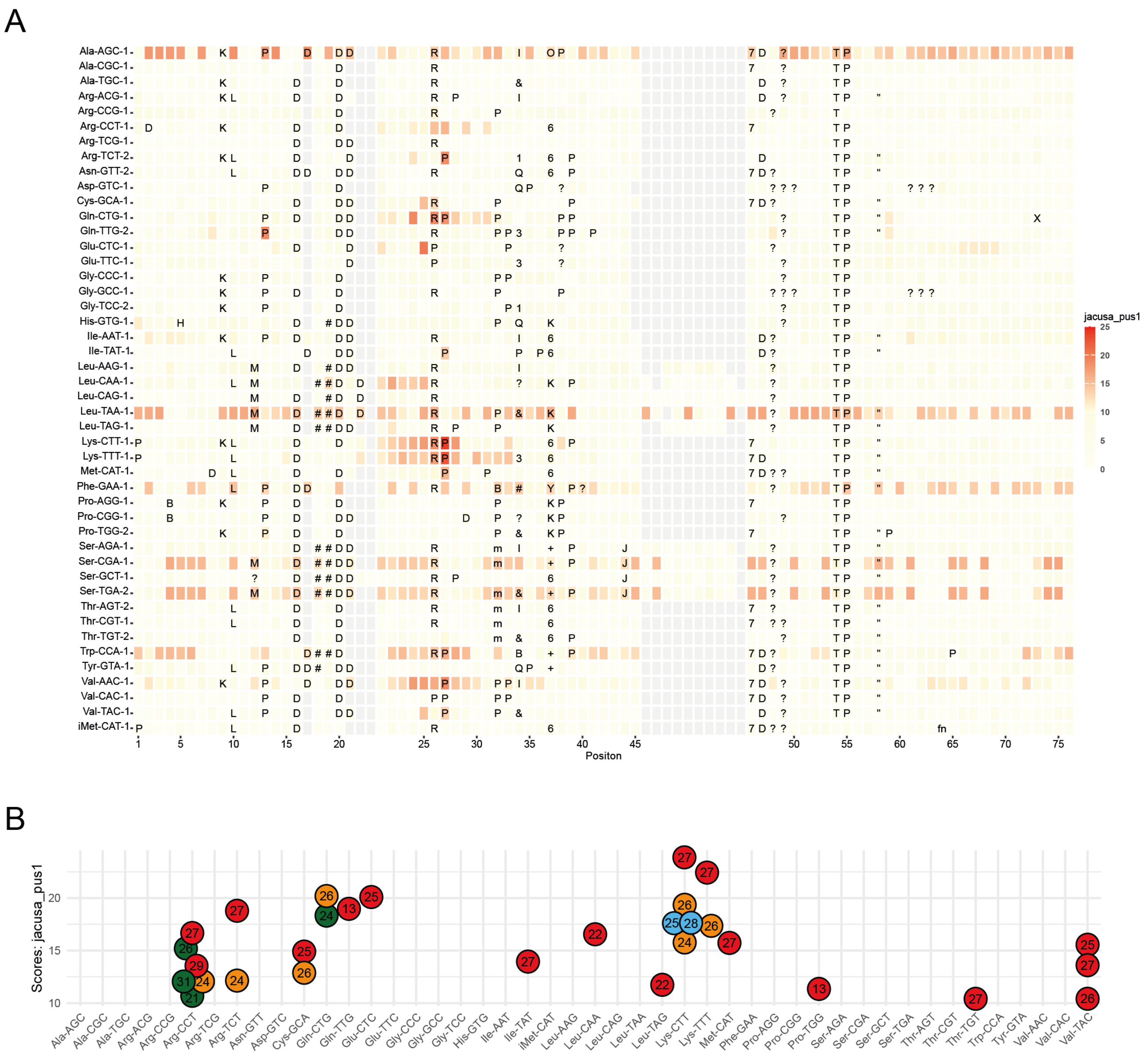


**Supplementary Figure 4**: Detection of Pus1-dependent tRNA modifications in *S. pombe*. A) Difference in basecalling errors between wildtype and *pus1Δ* as determined by JACUSA2. B) Filtered basecalling errors (JACUSA2 outlier values, filtered for outliers using Tukey's Fence (k=1.5, see Material and Methods, applied for each tRNA), values ≥ 10. Colors represent the reference nucleotide at the given position: U = red, A = green, C = blue, G = orange. Overlapping values were slightly shifted to improve visibility.

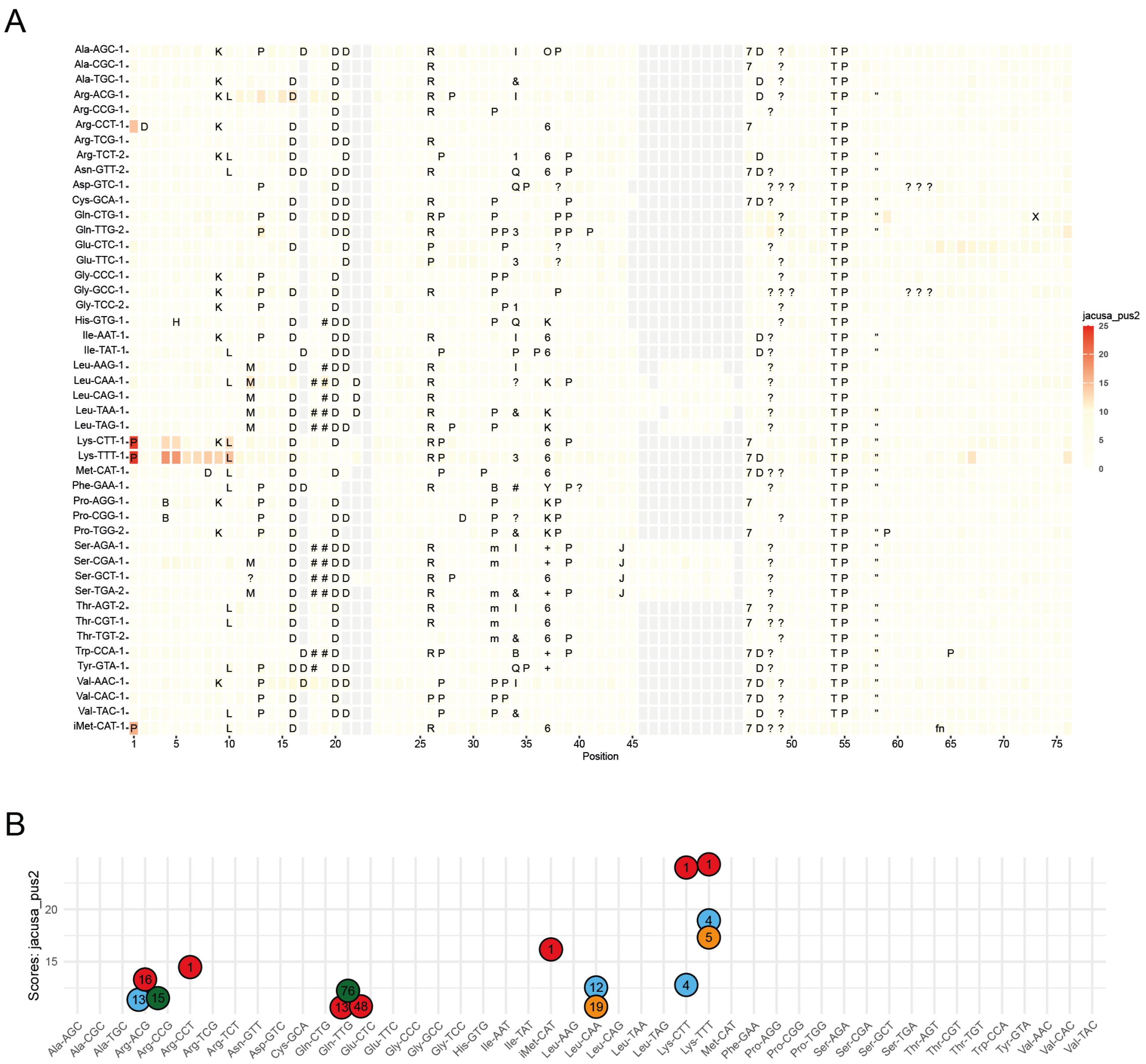


**Supplementary Figure 5**: Detection of Pus2-dependent tRNA modifications in *S. pombe*. A) Difference in basecalling errors between wildtype and *pus2Δ,* determined as in Suppl. Fig. 4. B) Filtered basecalling errors for wt/ *pus2Δ*, as determined in Suppl Fig. 4. Colours represent the reference nucleotide at the given position: U = red, A = green, C = blue, G = orange. Overlapping values were slightly shifted to increase visibility.

A

B

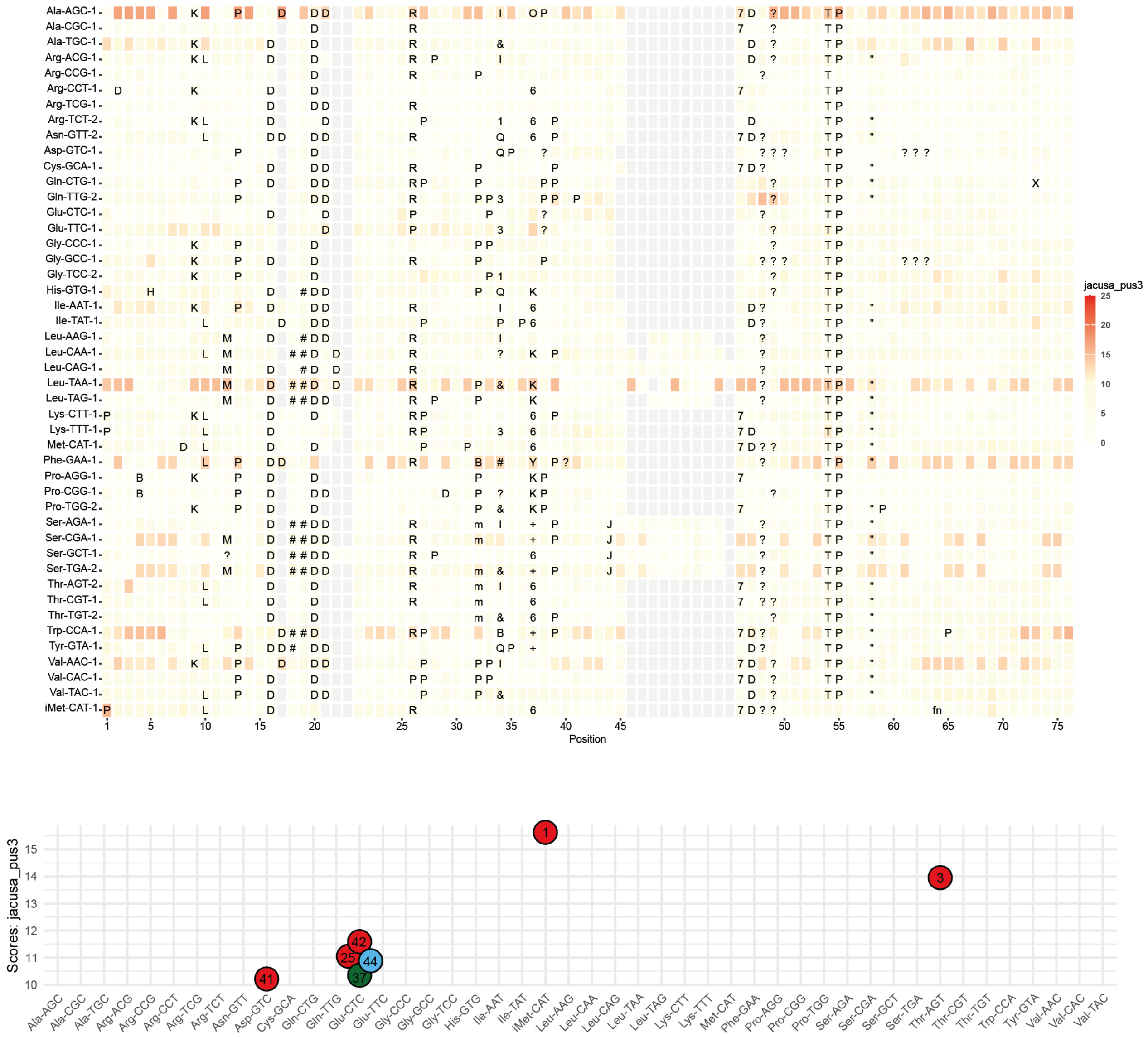


**Supplementary Figure 6**: Detection of Pus3-dependent tRNA modifications in *S. pombe*. A) Difference in basecalling errors between wildtype and *pus3Δ* strain, determined as in Suppl. Fig. 4. B) Filtered basecalling errors for wt/ *pus3Δ*, as determined in Suppl Fig. 4. Colors represent the reference nucleotide at the given position: U = red, A = green, C = blue, G = orange. Overlapping values were slightly shifted to increase visibility.

A

B

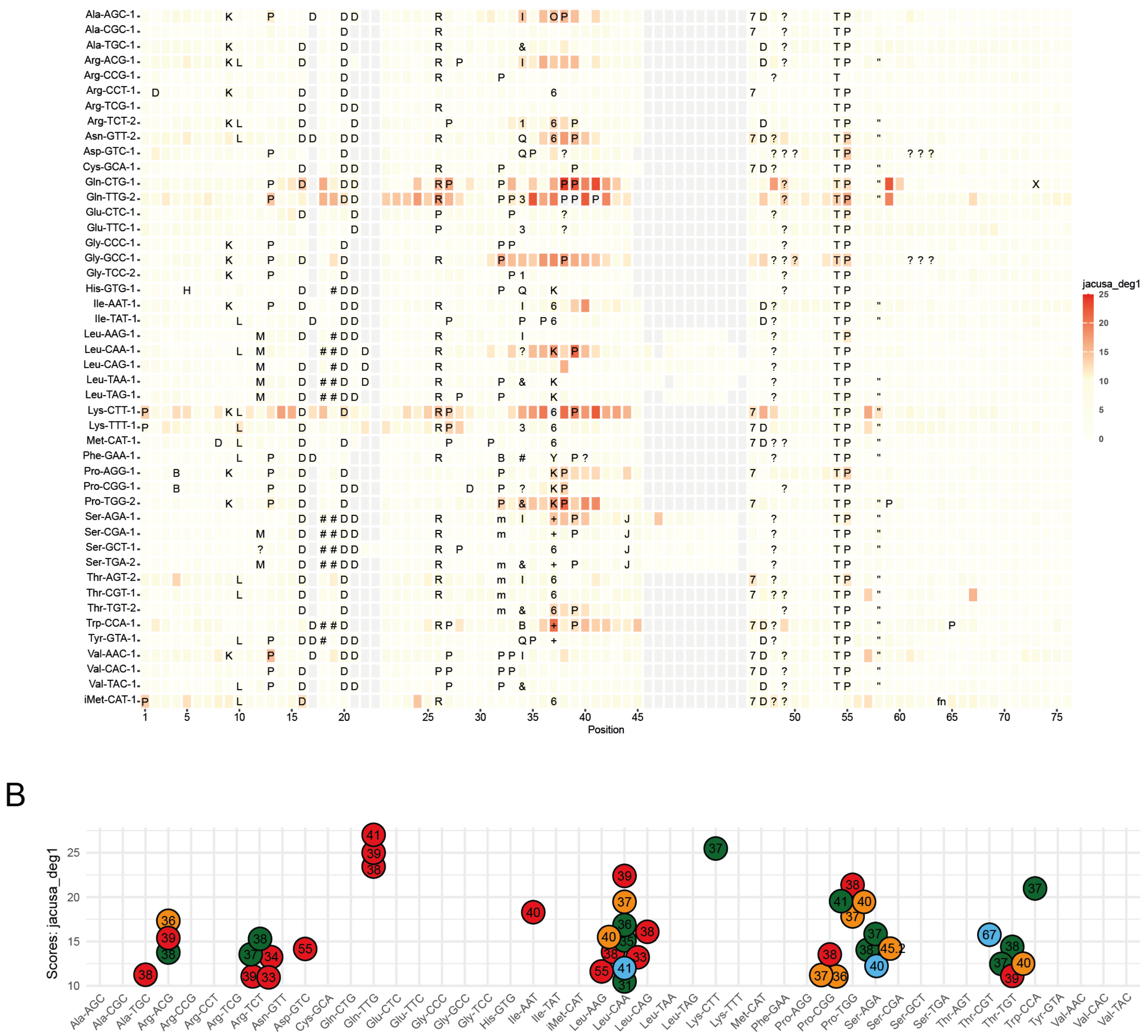

**Supplementary Figure 7**: Detection of Deg1-dependent tRNA modifications in *S. pombe*. A) Difference in basecalling errors between wildtype and *deg1Δ* strain, determined as in Suppl. Fig. 4. B) Filtered basecalling errors for wt/ *deg1Δ*, as determined in Suppl Fig. 4. Colors represent the reference nucleotide at the given position: U = red, A = green, C = blue, G = orange. Overlapping values were slightly shifted to increase visibility.

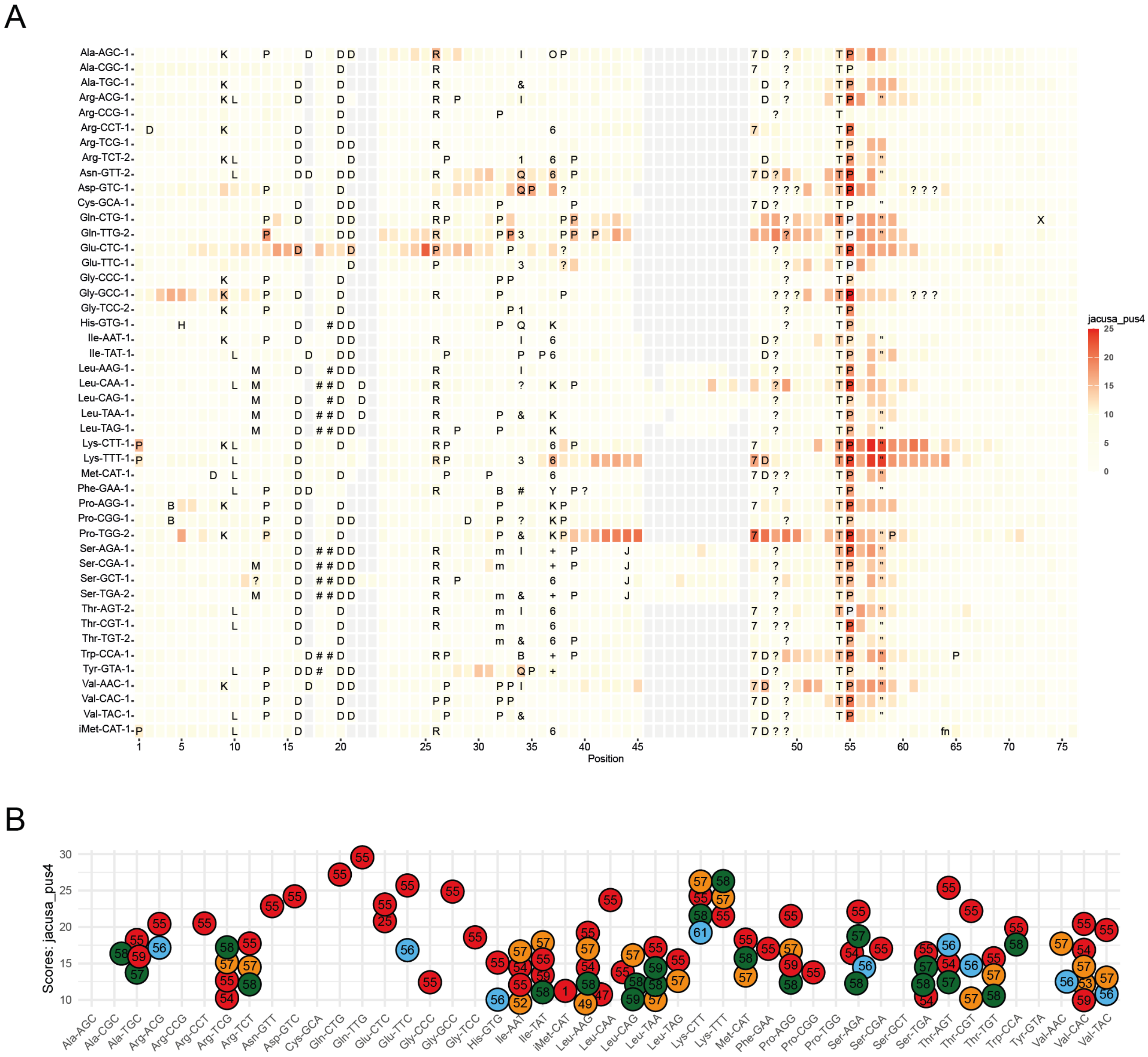


**Supplementary Figure 8**: Detection of Pus4-dependent tRNA modifications in *S. pombe*. A) Difference in basecalling errors between wildtype and *pus4Δ,* determined as in Suppl. Fig. 4. B) Filtered basecalling errors for wt/ *pus4Δ*, as determined in Suppl Fig. 4. Colours represent the reference nucleotide at the given position: U = red, A = green, C = blue, G = orange. Overlapping values were slightly shifted to increase visibility.

A

B

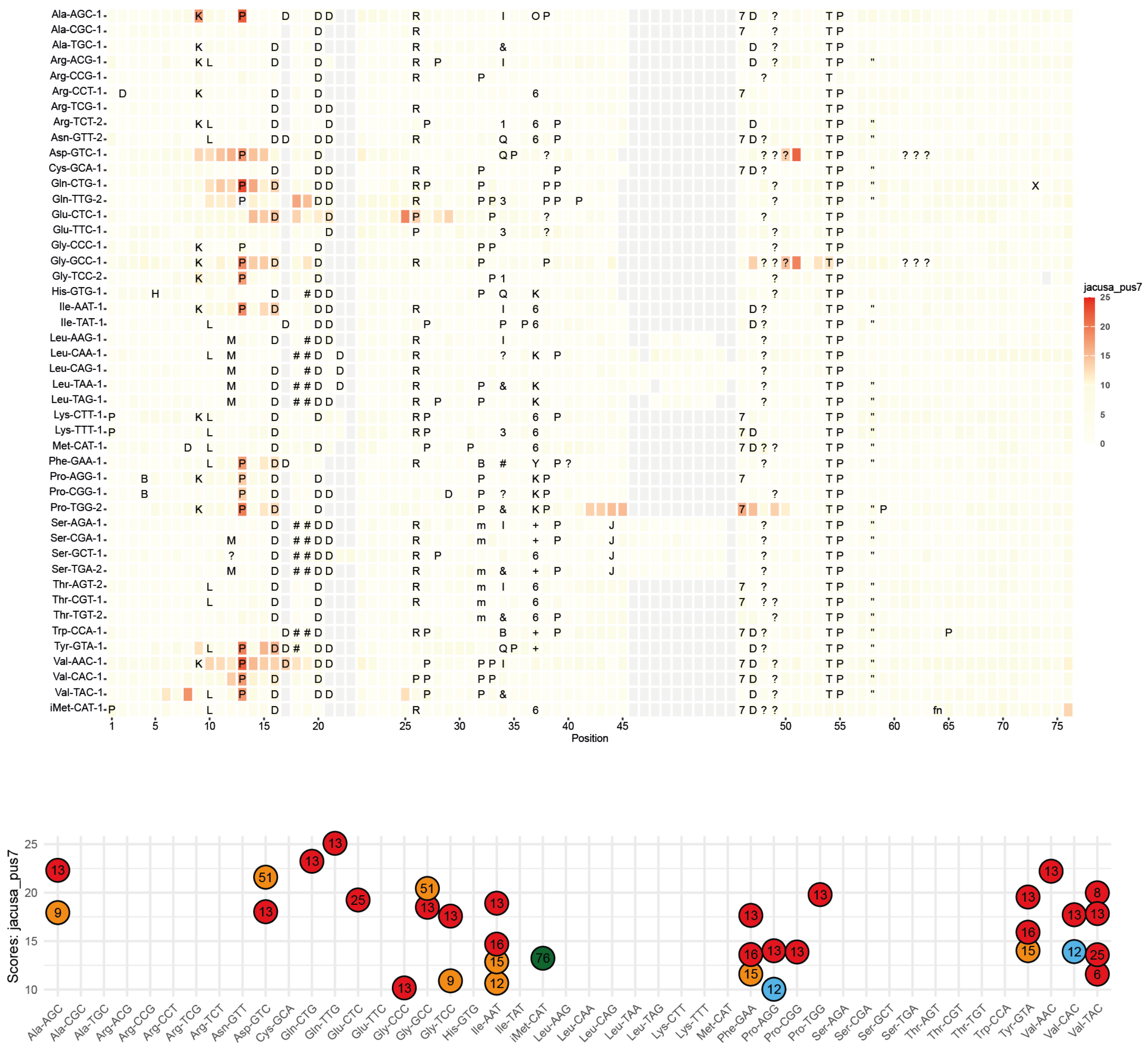

**Supplementary Figure 9**: Detection of Pus7-dependent tRNA modifications in *S. pombe*. A) Difference in basecalling errors between wildtype and *pus7Δ* determined as in Suppl. Fig. 4. B) Filtered basecalling errors for wt/ *pus7Δ*, as determined in Suppl Fig. 4. Colours represent the reference nucleotide at the given position: U = red, A = green, C = blue, G = orange. Overlapping values were slightly shifted to increase visibility.

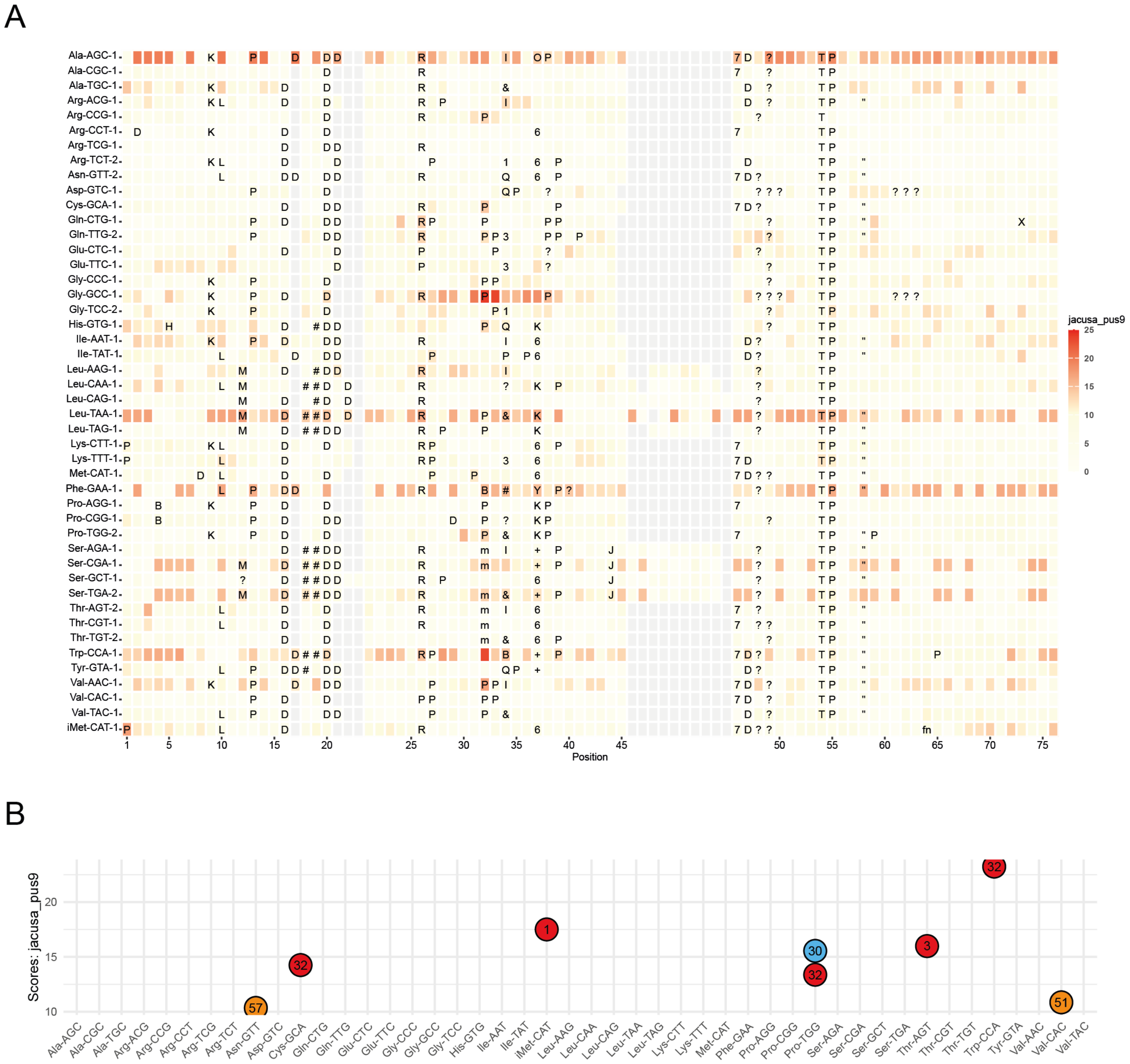


**Supplementary Figure 10**: Detection of Pus9-dependent tRNA modifications in *S. pombe*. A) Difference in basecalling errors between wildtype and *pus9Δ,* determined as in Suppl. Fig. 4. B) Filtered basecalling errors for wt/ *pus2Δ*, as determined in Suppl Fig. 4. Colours represent the reference nucleotide at the given position: U = red, A = green, C = blue, G = orange. Overlapping values were slightly shifted to increase visibility.

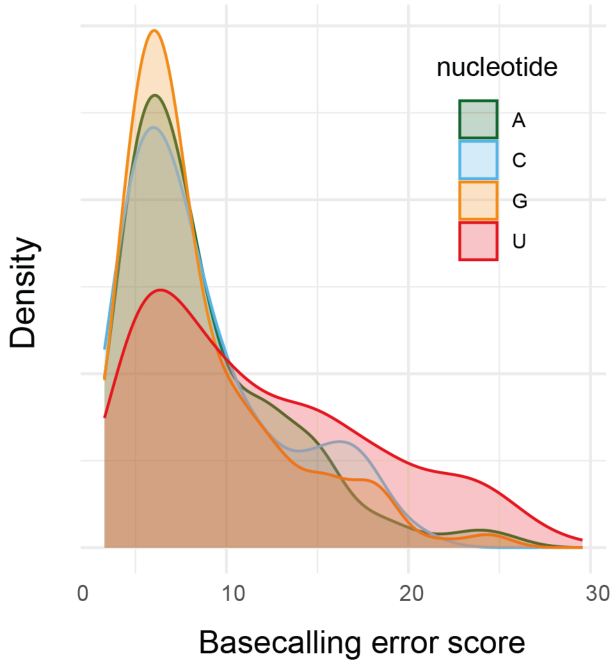


**Supplementary Figure 11**: Distribution of basecalling error outliers as defined by Tukey's Fence method. Colour gives the nucleotide identity as indicated.

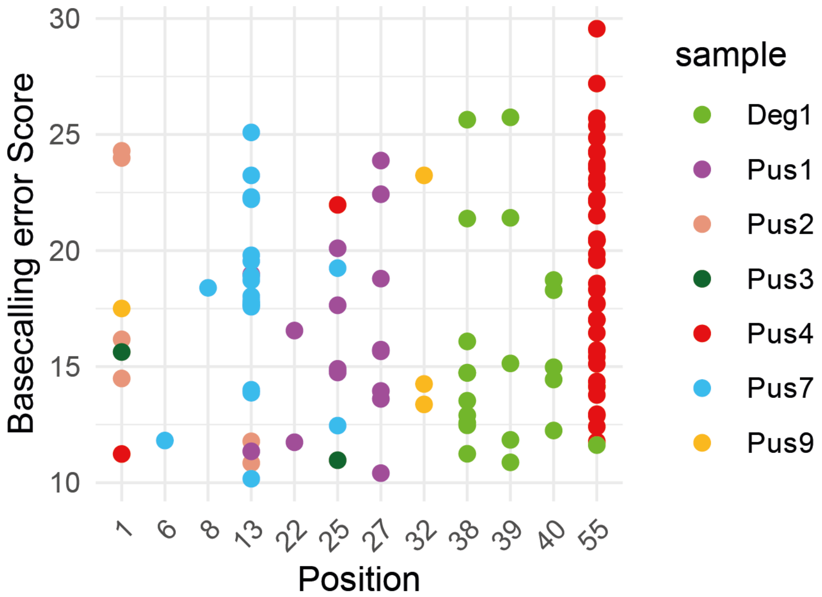


**Supplementary Figure 12**: Summary of basecalling error scores at U positions in direct sequencing of tRNAs from *S. pombe* strains lacking pseudouridine synthetases. The error scores indicate the position modified by the respective enzyme. Error scores are shown by tRNA position. Colours indicate the respective enzymes.

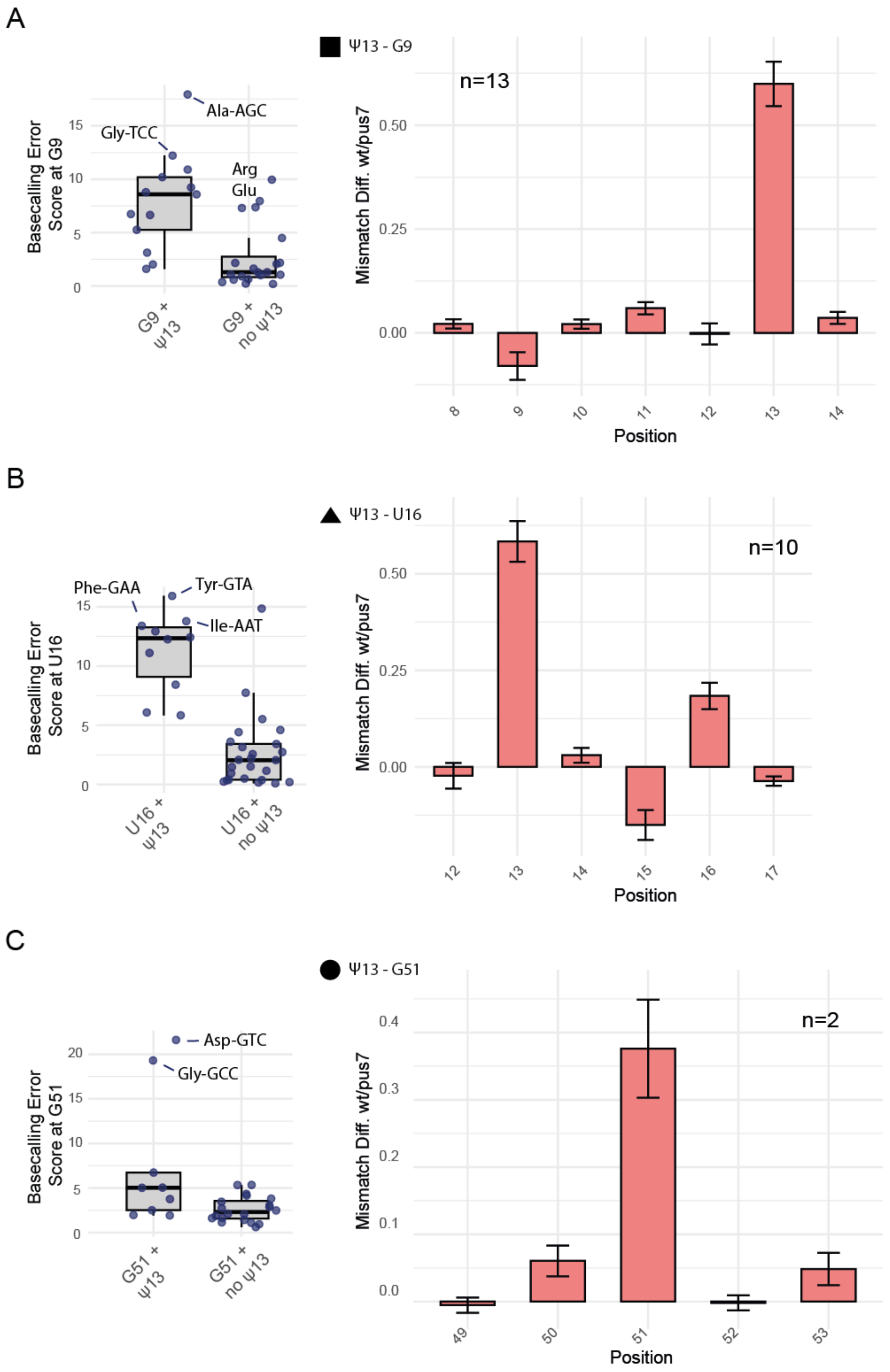


**Supplementary Figure 13**: Evaluation of potential Pus7-dependent tRNA modification crosstalk. A) Indication of crosstalk between Pus7-dependent Ψ13 and $m^1$G9. Left, basecalling error scores of wild-type (wt)/ *pus7Δ* at position G9 in tRNAs that also carry U13 compared to tRNAs with G9, but no U13. Error bars show standard deviation. Dots show the values of all tRNAs evaluated. Right, absolute difference in mismatch errors of wild-type (wt) compared to *pus7Δ* at positions 8 to 14. At position 13, wt shows more mismatch errors than *pus7Δ*, indicating more modification in wt, whereas there are less mismatch errors at position 9 in wt compared to *pus7Δ*, suggesting that position G9 is less modified in wt than *pus7Δ*. n represents the total number of tRNAs with G9 and Ψ13 for which the mismatch differences were calculated. B) as in A, but evaluating positions U13 and U16, n represents the total number of tRNAs with U16 and Ψ13 for which the mismatch differences were calculated, left part as shown in Figure 3E. C) as in A, but evaluating positions U13 and G51, n represents the two outlier tRNAs of the G51 and Ψ13 group for which the mismatch differences were calculated.

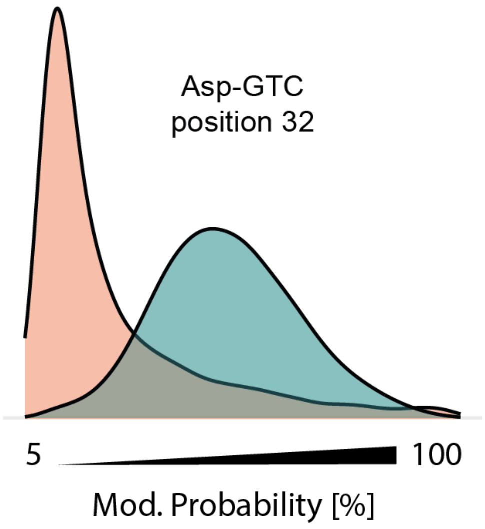


**Supplementary Figure 14**: The Dorado Ψ caller senses Q34 modification. Ψ modification probability at U32 of tRNA-Asp, which lies 5' to G34/ Q34. Values of Ψ calling for wt cells cultured with (Q+) or without (Q-) queuine are shown. In the presence of Q modification, the Dorado Ψ caller erroneously predicts Ψ32.

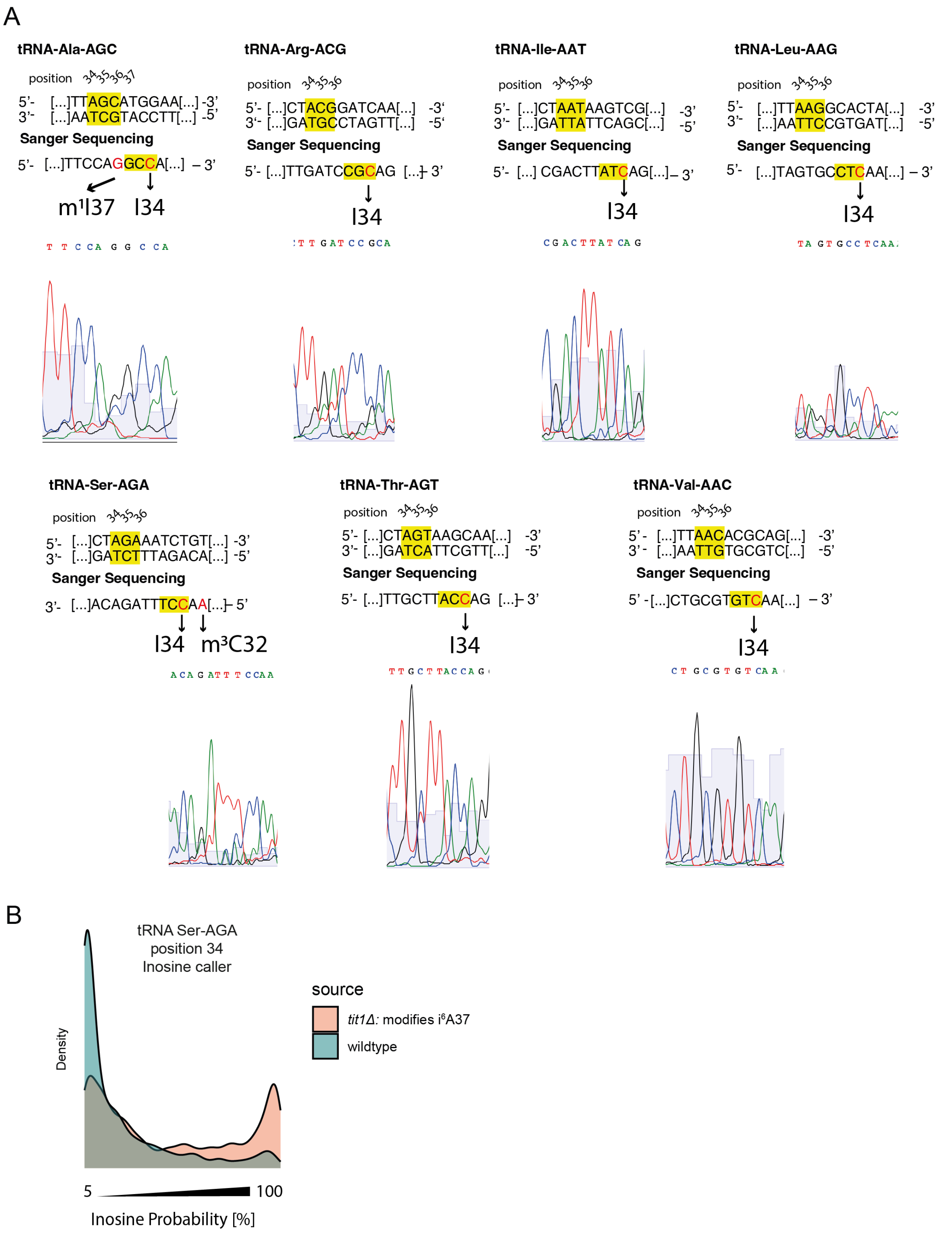


**Supplementary Figure 15**: A) Detection of inosine modification in tRNAs from *S. pombe* by reverse transcription, PCR amplification and Sanger sequencing. Yellow shows the anticodon, red shows the characteristic point mutations introduced after reverse transcription, amplification and sequencing. Inosine can be identified by the characteristic A to G change (visible as C in the sequence of the reverse strand). Sequencing chromatograms of the relevant sequence regions are shown. For Ala-AGC, the methylated inosine at position 37 can be identified by A to C change (visible as G in the sequencing result). For Ser-AGA, m³C at position 32 results in a mismatch. B) Inosine 34 probability in Ser-AGA as determined by the inosine caller in wild-type and *tit1Δ*.

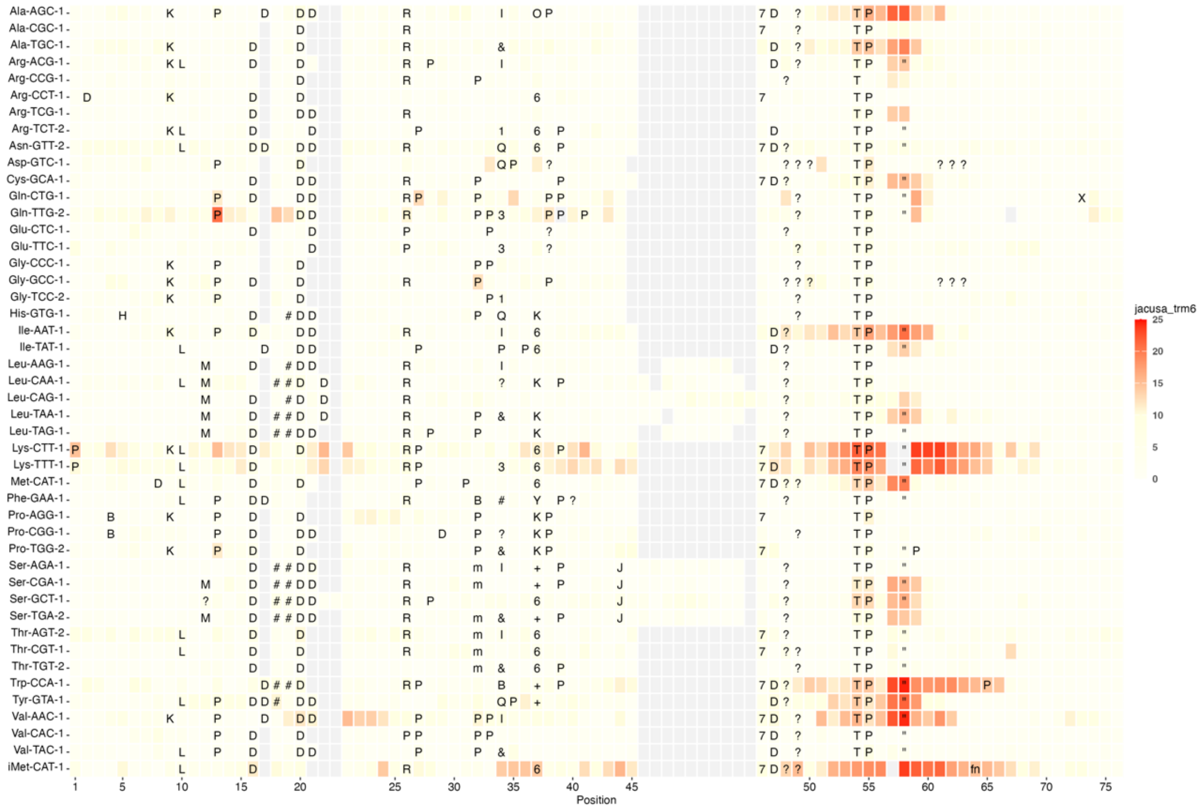


**Supplementary Figure 16**: Detection of Trm6-dependent m¹A modifications of tRNA in *S. pombe*. Difference in basecalling errors between wild-type and *trm6*Δ, determined as in Suppl. Fig. 4.

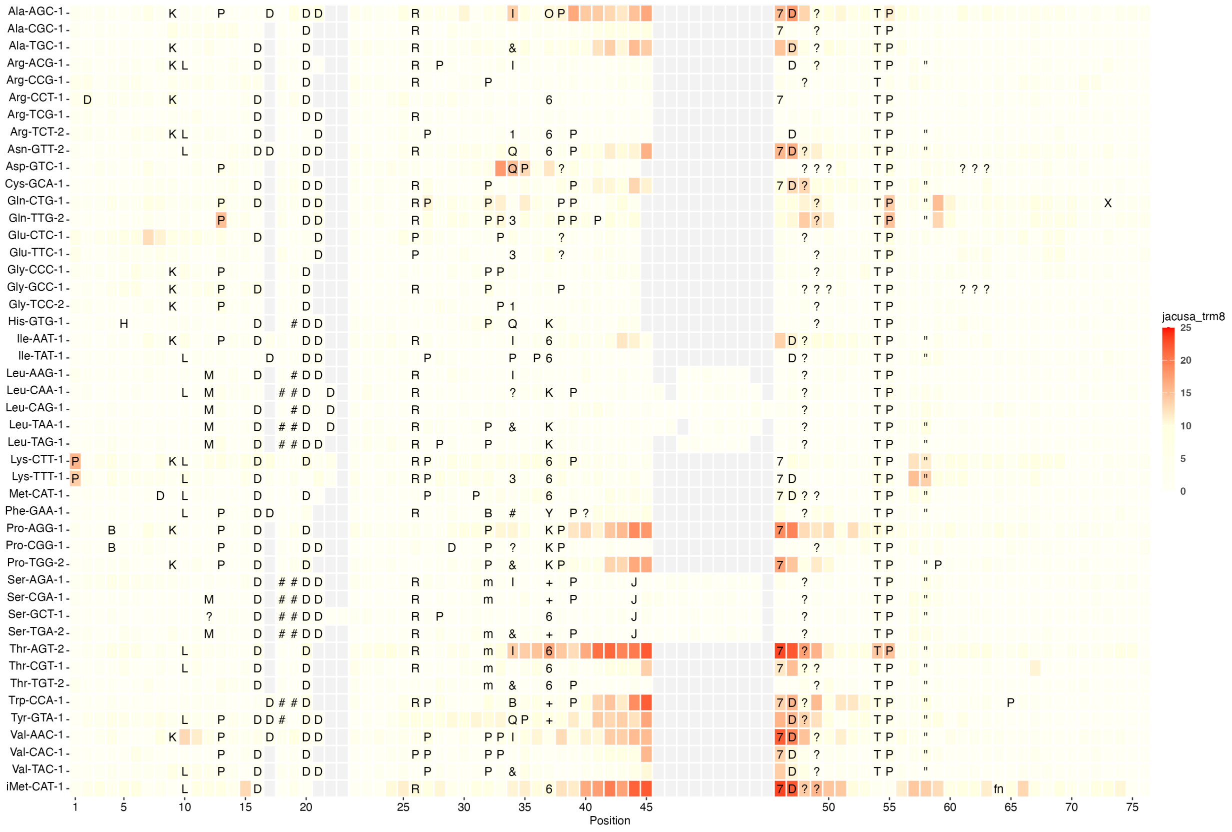


**Supplementary Figure 17**: Detection of Trm8-dependent m⁷G modification on tRNA in *S. pombe*. Difference in basecalling errors between wild-type and *trm8*Δ, determined as in Suppl. Fig. 4.

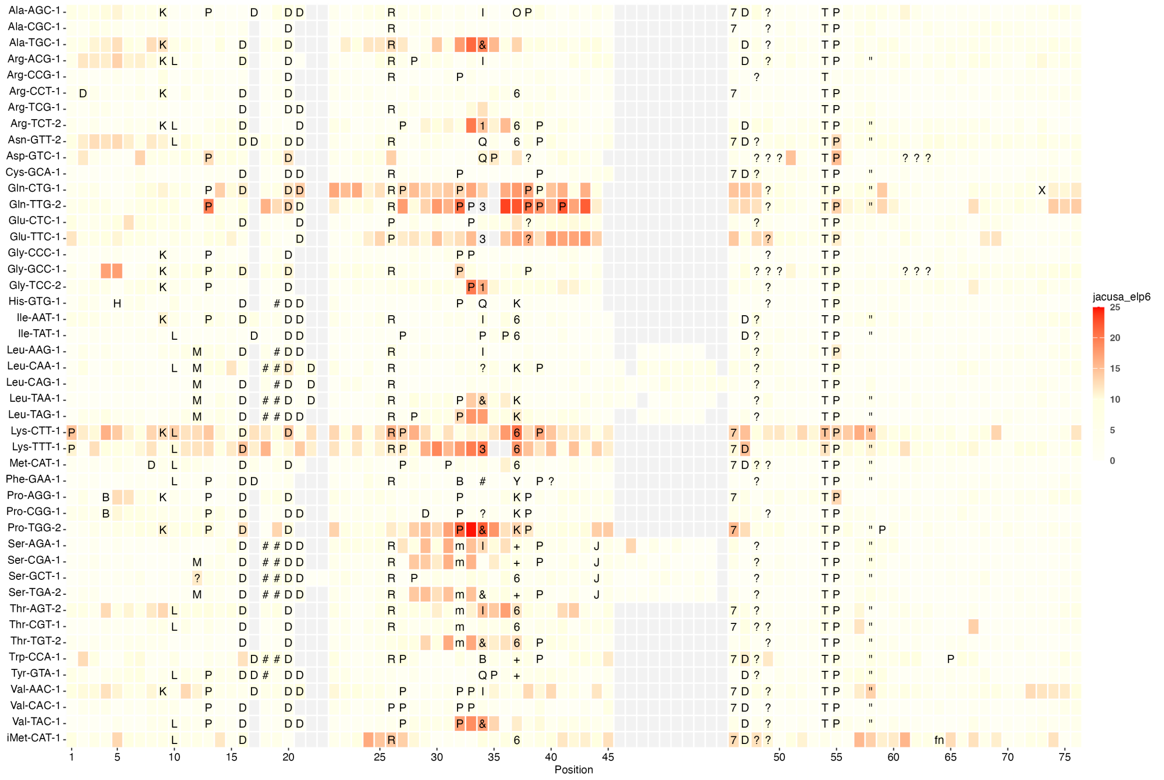

**Supplementary Figure 18**: Detection of Elp6-dependent U34 modification on tRNA in *S. pombe*. Difference in basecalling errors between wild-type and *elp6Δ*, determined as in Suppl. Fig. 4.

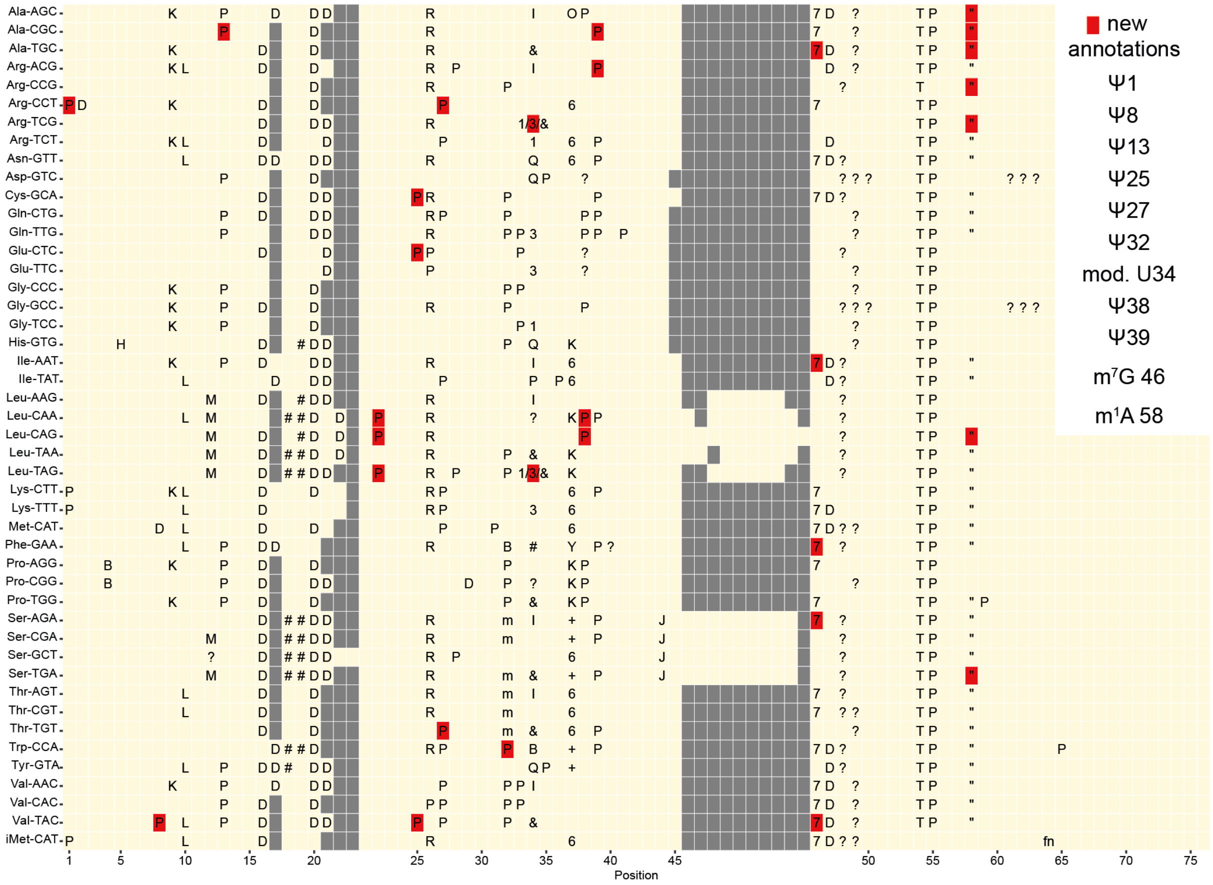

**Supplementary Figure 19**: Novel tRNA modifications (Ψ, m¹A, m⁷G, red) detected by dRNA-seq in this study. One-letter-code as in Supplementary Figure 1.
